# SIV infection disrupts the spatial cellular and communication networks of pulmonary granulomas during SIV/Mtb co-infection

**DOI:** 10.64898/2026.04.24.720743

**Authors:** Jessica M. Medrano, Persis Sunny, Collin R. Diedrich, Pauline Maiello, Christopher Kline, Tara Rutledge, Edwin Klein, Joshua Mattila, Jishnu Das, Zandrea Ambrose, Philana Ling Lin

## Abstract

Tuberculosis (TB) caused by *Mycobacterium tuberculosis* (Mtb) is the leading infectious cause of death globally. Despite wide use of antiretroviral therapy (ART) by people living with HIV, the risk of TB remains increased. To understand immune interactions within lung granulomas, we compared spatial transcriptomics of Mtb and simian immunodeficiency virus (SIV) in co-infected macaques with or without ART as a model for HIV/Mtb co-infection. Spatially differentiated transcriptional profiles were observed in Mtb-only granulomas, with myeloid cells enriched in metabolic/antimicrobial pathways within the inner ring and T cell costimulatory/activation pathways enriched in the outer ring. These spatially distinct patterns were lost in SIV/Mtb granulomas with higher enrichment in type I IFN pathways compared to Mtb-only granulomas. SIV/ART/Mtb granulomas had an intermediate transcriptional pattern without restoration to Mtb-only granulomas, despite a lack of viral replication. Cell-cell communication was reduced among SIV/Mtb co-infected groups. These data suggest that HIV disrupts spatially organized immune functions of granulomas, which are not fully restored by ART.

## Introduction

Infection with *Mycobacterium tuberculosis* (Mtb) continues to be the leading cause of death from a single infectious agent, with over 10.7 million new cases of TB and 1.23 million deaths in 2024 and an estimated a one-quarter of the world having been exposed to the bacteria (1). About 10% of those infected with Mtb will go on to develop active disease, typically within the first two years post-infection (2). While the immunologic mechanisms necessary to control Mtb infection are not fully understood, the most important risk factor for severe TB disease is co-infection with human immunodeficiency virus (HIV) (1). The resurgence of the tuberculosis (TB) epidemic is closely tied to the emergence of HIV, and disease burden is concentrated in overlapping populations in southern Africa (1). These two pathogens have a synergistic relationship where each drives worse outcomes of the other: TB is the leading cause of death among people living with HIV (PLWH) with one-third of HIV-associated deaths are from TB (1, 3). About 73% of people living with HIV (PLWH) are on antiretroviral therapy (ART), which, while not curative, prevents ongoing HIV replication and restores CD4+ T cells. However, PLWH are still more likely to develop TB, despite suppressive ART (4).

The hallmark of TB is the granuloma: an organized collection of immune cells often with a necrotic central area of Mtb-infected macrophages surrounded by a ring of lymphocytes and other innate immune cells (reviewed in (5, 6)). Granulomas form within the lung parenchyma but can also spread to extrapulmonary sites including lymph nodes, spleen, liver, central nervous system, and gastrointestinal/genitourinary tracts (reviewed in (7)). The granuloma is the primary site of immune interactions with Mtb. From the host’s perspective, granulomas function to contain Mtb, but Mtb has also evolved to persist within them (8). Their structure is heterogeneous both in the spectrum of granuloma types (e.g., mineralized, non-necrotic, and necrotic granulomas) and in the complex populations of macrophages and other immune cells present within a granuloma (5, 9, 10). Each granuloma is independent within the same host and even within the same lung lobe, and the clinical outcome of Mtb infection depends on the collective success or failure of multiple granulomas in inhibiting Mtb growth (10).

How HIV disrupts the protective immune responses against Mtb within granulomas remains incompletely understood, though studies with simian immunodeficiency virus (SIV) in non-human primates (NHP) have shown that cells within the granuloma can be infected with SIV during co-infection (11–13). CD4+ T cells are the most influential adaptive immune cell type in mediating protection from Mtb, and are critical for activating macrophages, the first cell type to encounter Mtb, through secretion of cytokines produced by CD4+ T cells, including IFN-γ and TNF (14). In vitro, HIV infection of human peripheral blood mononuclear cells (PBMCs) increases Mtb burden in monocyte-derived macrophages (15); and in vivo, CD4+ T cells within the granuloma are depleted as early as two weeks post-SIV infection (12). Depletion of CD4+ T cells during HIV or SIV co-infection is correlated with worse TB outcomes (4, 12, 16). Importantly, we and others have shown that SIV-induced reactivation of latent Mtb infection can occur independent of CD4+ T cells loss (11, 17). Work in NHPs has demonstrated that immune-driving enzyme and cytokine protein expression in Mtb granulomas is spatially organized (9, 18–20), though the mechanisms by which T cell and macrophage communication in granulomas is altered during SIV co-infection is not fully elucidated.

Here, we used an NHP model of SIV and subsequent Mtb co-infection with and without suppressive ART to interrogate the immune cell dynamics in granulomas. During Mtb infection, cellular functions within the granuloma were spatially distinct, such that myeloid and T cell gene expression profiles were different within the inner and outer regions. Co-infection with SIV, which resulted in higher bacterial burden in the granuloma and severe TB disease, led to loss of functional distinction between the inner and outer regions, for both myeloid cells and T cells, and a reduction in overall cell-cell communication within the granuloma primarily driven by a reduction in T cell signaling. The reduction in functional organization of cells in granulomas was not fully restored in SIV/Mtb co-infected animals treated with ART, highlighting that while suppression of SIV viremia reduced bacterial burden and improved outcomes, ART did not rescue all aspects of SIV/Mtb pathogenesis (16, 21). Understanding how HIV alters dynamics within granulomas and drives increased TB disease during co-infection is critical for vaccine and therapeutics development.

## Results

### Myeloid and T cell function in granulomas are altered during SIV/Mtb co-infection

To understand the effect of HIV on cellular dynamics in Mtb granulomas, we used an NHP model of HIV/Mtb co-infection, using SIV as a surrogate for HIV. Co-infected animals were infected with pathogenic SIV with or without daily ART starting at day 3 post-infection followed by low-dose Mtb challenge 16 weeks later (SIV/Mtb and SIV/ART/Mtb, respectively). [Figure 1A, (16)]. A control group of animals was infected with Mtb only. Plasma viremia peaked one week after infection before establishing a set-point in SIV/Mtb animals, and all ART-treated animals had undetectable plasma viremia by 4 weeks post-infection. CD4+ T cells were depleted in the blood and lung granulomas in SIV/Mtb but not SIV/ART/Mtb or Mtb animals (16). Mtb bacterial burden per granuloma was higher in SIV/Mtb granulomas compared to both Mtb only and SIV/ART/Mtb animals [Figure 1B, (16)]. Almost all granulomas from SIV/Mtb animals had detectable SIV RNA by qRT-PCR (Figure 1C), whereas a selection of granulomas from SIV/ART/Mtb granulomas did not have detectable SIV RNA, as expected (16). Despite higher bacterial burden, SIV/Mtb granulomas were not appreciably different histologically from Mtb or SIV/ART/Mtb granulomas when assessed via H&E staining (Supplemental Figure 1)(13)￼. Thus, we hypothesized that the differences in SIV/Mtb granulomas may be related to cellular function rather than overall granuloma architecture.

**Figure 1.**
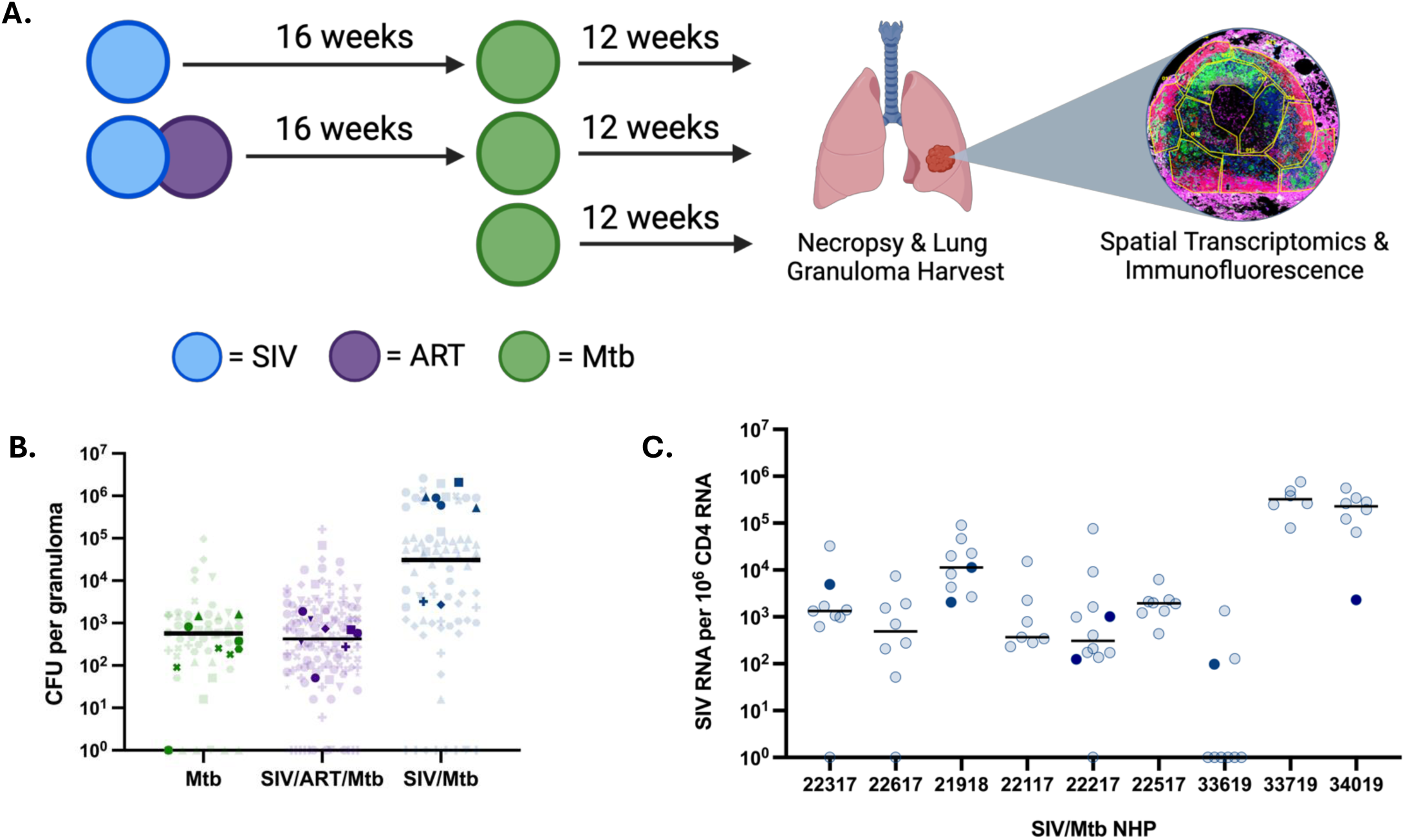
Study design and granuloma-specific Mtb and SIV quantitation. (A) Macaques were infected with SIV (16 weeks) and Mtb (12 weeks) [top row, n = 9], SIV (16 weeks) and Mtb (12 weeks) with ART beginning 3 days post-SIV infection [middle row, n = 10], or Mtb only (3 months) [bottom row, n = 9). Macaques were necropsied and individual lung granulomas were harvested and assessed for Mtb bacterial colony forming units (CFU), SIV RNA copies, and histology. A subset of granulomas was assessed for spatial transcriptomics (Nanostring GeoMX) and immunofluorescence. (B) Total Mtb CFU in each granuloma isolated at necropsy. Each symbol is a granuloma; each shape is an NHP. Horizontal lines indicate medians per group. All granulomas harvested are shown with dark shapes indicating the granulomas used for GeoMX. Data were adapted from (Diedrich et al., 2025). (C) SIV RNA copies per 10^6^ CD4 RNA copies in lung granulomas at necropsy. Each circle is a granuloma; horizontal lines indicate medians for each animal. All granulomas harvested are shown with dark symbols indicating the granulomas used for GeoMX.

To assess the impact of SIV infection on immunological responses within the granuloma, spatially defined transcriptomic changes in granulomas were assessed using the Nanostring GeoMX platform (Supplemental Table 1). In this assay, barcoded RNA probes targeting the whole transcriptome and fluorescent antibodies for T cells (CD3) and myeloid cells (CD68) were used. Regions of interest (ROIs) of the inner and outer rings (excluding the necrotic, acellular core) of the Mtb granulomas were manually delineated using serial hematoxylin and eosin staining. The transcriptome of CD68+ myeloid cells and CD3+ T cells from each region of lung granuloma were interrogated (Supplemental Figure 2).

Differentially expressed genes (DEG) from myeloid and T cells within entire granulomas (regardless of inner or outer ring) were compared between groups. Expression patterns in SIV/Mtb myeloid cells were distinct from both the transcriptome in Mtb only myeloid cells (1205 differentially expressed genes; Supplemental Table 2) and SIV/ART/Mtb myeloid cells (1418 differentially expressed genes; Supplemental Table 2, Figure 2A). Gene set enrichment analysis (GSEA) was performed to compare pathways enriched in DEGs in myeloid cells from each group. Pro-inflammatory pathways were the most dramatically enriched in DEGs from myeloid cells in SIV/Mtb granulomas compared to both the SIV/ART/Mtb and Mtb only granulomas (Figure 2B, Supplemental Figure 3, Supplemental Table 3). Myeloid cells from SIV/Mtb granulomas overall were significantly enriched in DEGS from type I interferon signaling pathways, acute inflammatory responses, IL-1 signaling, chemokine production, and response to chemokines compared to myeloid cells from Mtb and SIV/ART/Mtb granulomas (Figure 2C, Supplemental Table 3). Myeloid cells from the SIV/ART/Mtb granulomas differed in their enrichment pattern for DEGs in specific pathways: acute inflammatory responses (complement) and chemokine production DEGS were enriched relative to Mtb, though not identical to Mtb only granulomas (Figure 2C, Supplemental Table 3). However, IL-1 signaling and interferon alpha/beta signaling, though not enriched overall in SIV/ART/Mtb myeloid cells, still had distinct gene expression patterns from Mtb only myeloid cells (Figure 2C). Thus, myeloid cells contribute to a more inflammatory environment in SIV/Mtb granulomas, and ART can restore some, but not all, aspects of myeloid cell function.

**Figure 2.**
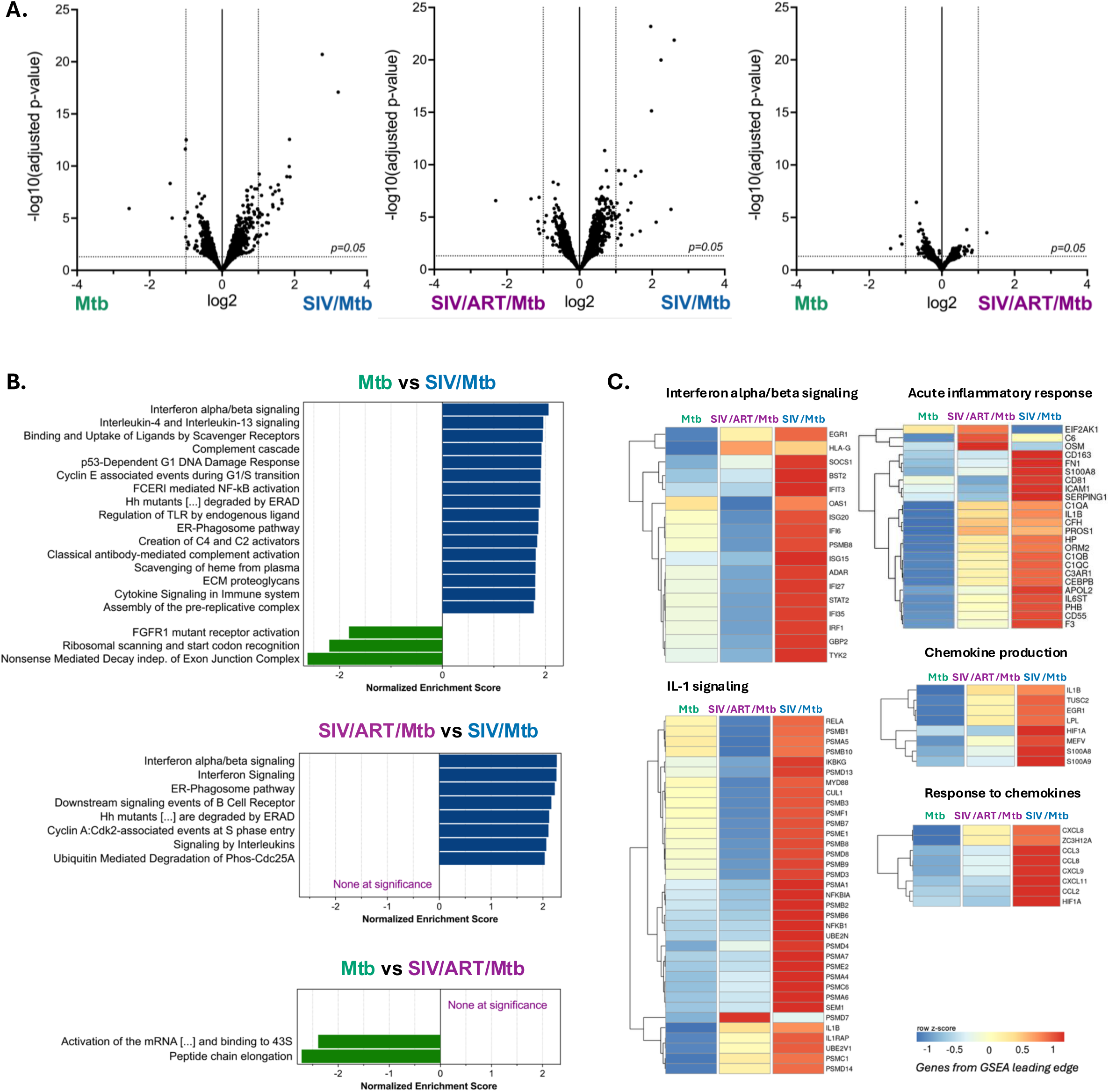
Myeloid cell function in granulomas is altered during SIV/Mtb co-infection. (A) Volcano plots of differential gene expression in myeloid cells in whole granulomas between groups. Each point represents one gene; horizontal dotted lines indicate adjusted p-values at 0.05 and vertical lines indicate 1 and -1 log-fold change. (B) Gene set enrichment analysis (reduced redundancy) of all myeloid cells in granulomas between groups. Pathways are from WebGestalt using the Reactome database. Only pathways with adjusted p-values < 0.5 are shown. (C) Z-scores of gene expression in genes from pro-inflammatory pathways in all myeloid cells from granulomas in each group. Genes are Z-scored by row.

Differentially expressed genes from all granuloma T cells were also compared between groups (Figure 3A). As with myeloid cells, the SIV/Mtb T cell transcriptome was distinct from both SIV/ART/Mtb T cells (1032 differentially expressed genes; Supplemental Table 4) and Mtb only T cells (1412 differentially expressed genes; Supplemental Table 4). SIV/ART/Mtb and Mtb only T cell transcriptomes were more similar in gene expression patterns to each other (116 differentially expressed genes; Supplemental Table 4,Figure 3A). Based on GSEA analysis, T cells from the SIV/Mtb group had a significant reduction in gene expression in pathways related to T cell activation and signaling, including co-stimulation by CD28, PD-1 signaling and IL-2 family signaling, compared to T cells in either the SIV/ART/Mtb and Mtb only groups (Figure 3B, Supplemental Figure 4, Supplemental Table 5). Enrichment of DEGS in these pathways in SIV/ART/Mtb and Mtb T cells was similar (Figure 3C, Supplemental Table 5). Thus, T cells in SIV/Mtb granulomas were overall less activated and had less downstream signaling than SIV/ART/Mtb and Mtb only T cells, which ART was able to restore to a similar level as T cells in Mtb only granulomas.

**Figure 3.**
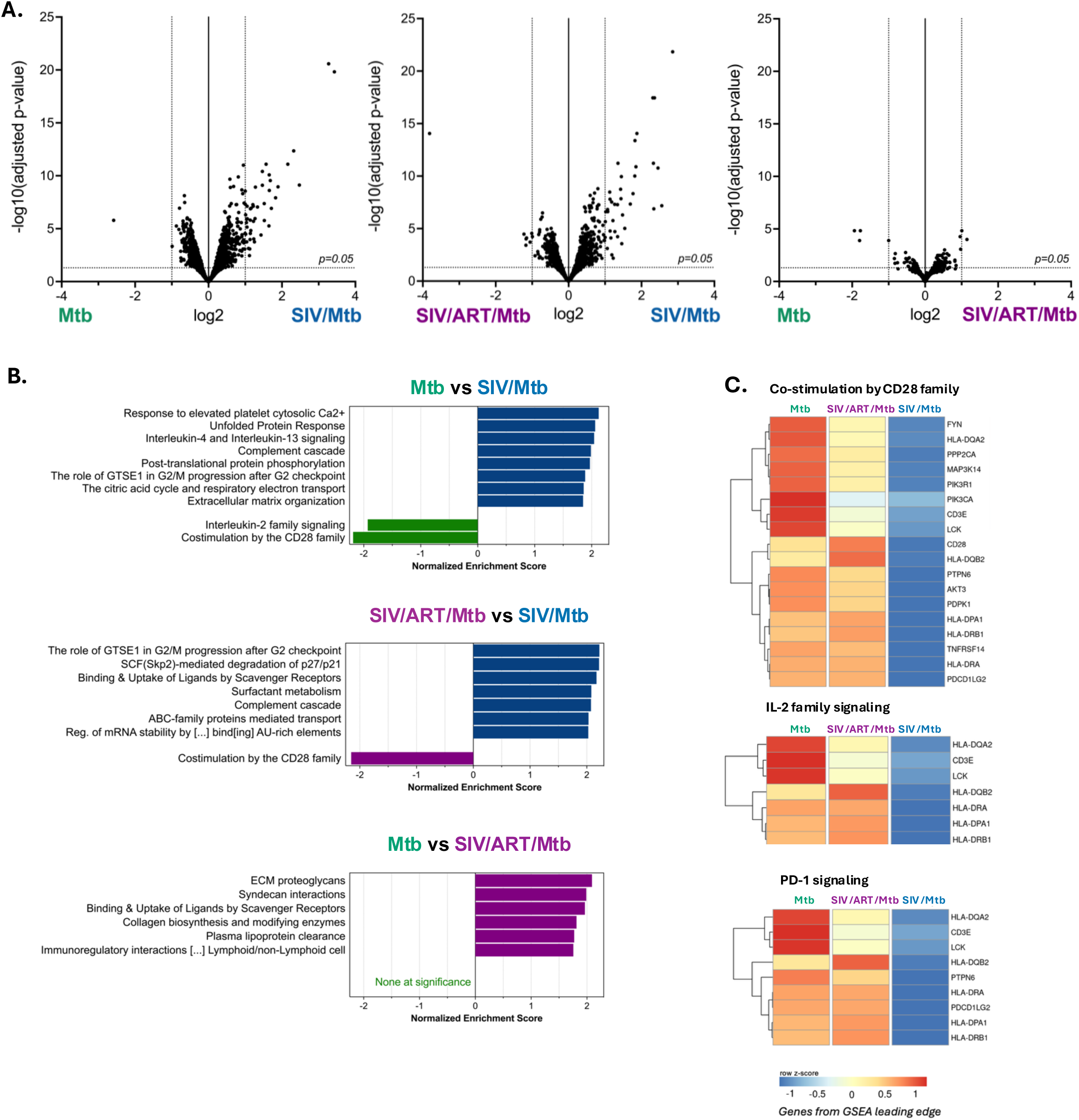
T cell function in granulomas is altered during SIV/Mtb-co-infection. (A) Volcano plot of differential gene expression in all T cells in granulomas between groups. Each point is one gene; horizontal dotted lines indicate adjusted p-values at 0.05 and vertical lines indicate 1 and -1 log-fold change. (B) GSEA (reduced redundancy) of all T cells in granulomas between groups. Pathways are from WebGestalt using the Reactome database. Only pathways with adjusted p-values < 0.5 are shown. (C) Z-score of gene expression in genes from T cell activation and signaling pathways in all T cells in granulomas in each group. Genes are Z-scored by row.

### Spatial distinction between the inner and outer portions of the granulomas among myeloid cell and T lymphocyte function is impaired during co-infection

As granulomas are highly structured environments, expression patterns of myeloid cell ROIs from the inner and outer rings of granulomas within each group were compared to understand how these transcriptional changes were spatially patterned. The transcriptomes of the inner and outer myeloid cells in Mtb only granulomas were distinct from each other, with 1067 differentially expressed genes in the inner region compared to the outer region (Figure 4A, Supplemental Table 6). Enriched DEGs in myeloid cells in the inner versus outer regions of these Mtb only granulomas were predominantly in pathways related to metabolism and antimicrobial function (Supplemental Table 7). In contrast, the transcriptomes of myeloid cells in SIV/Mtb granulomas in inner and outer regions were less distinct from each other (401 differentially expressed genes; Figure 4A, Supplemental Table 6), with a few unique pathways in the inner cellular ring (Supplemental Table 7). Spatial analysis of gene expression in SIV/ART/Mtb myeloid cells revealed that gene expression in the inner and outer rings were not as spatially distinct as tin the Mtb only group (293 differentially expressed genes; Figure 4A, Supplemental Table 7), although the DEGs were enriched in pathways also present in Mtb granulomas (Supplemental Table 7). Thus, while ART can restore some overall cellular functions in granulomas, it does not fully restore the spatial arrangement of their functions.

**Figure 4.**
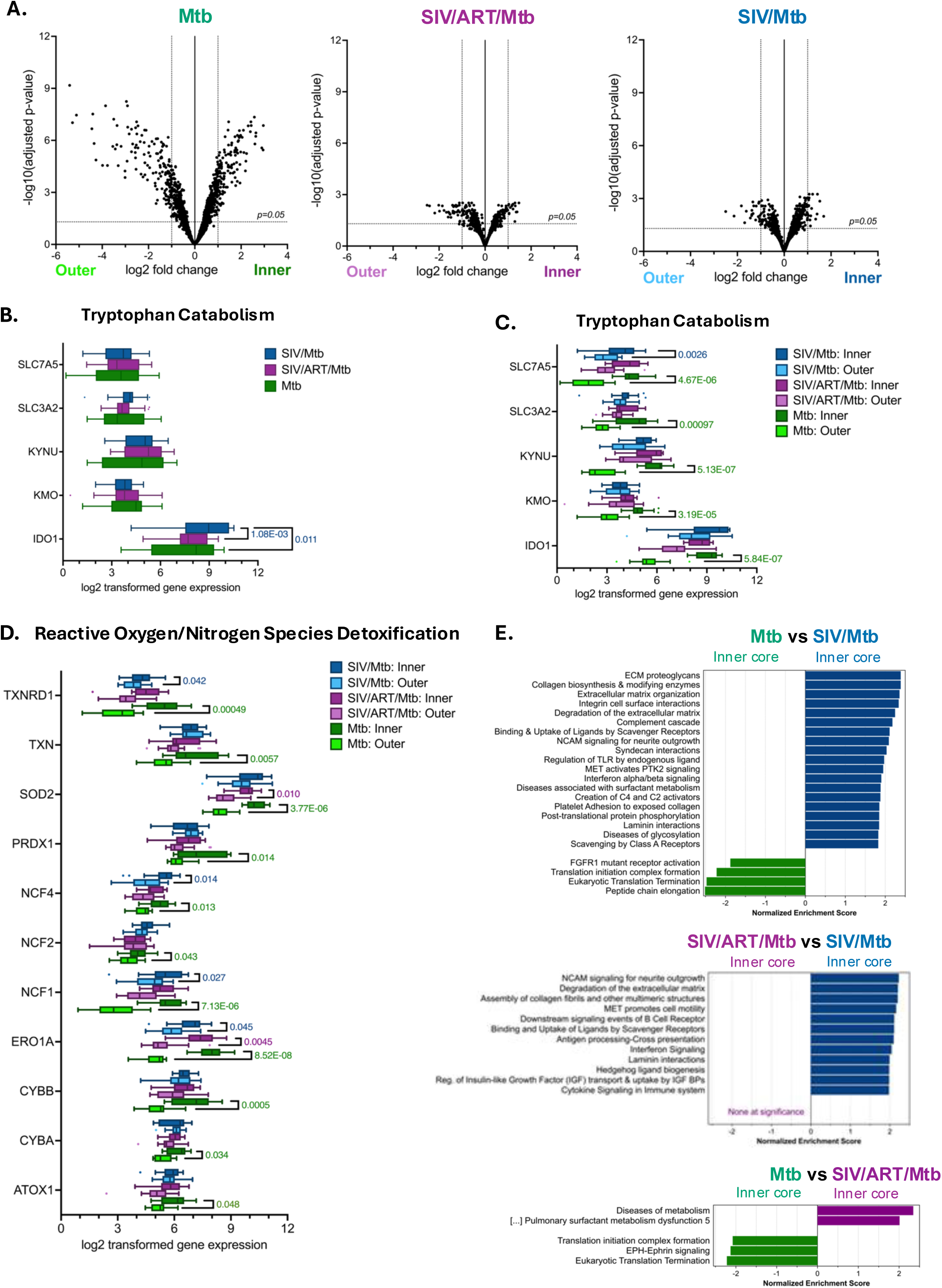
Spatial organization of myeloid cell function in granulomas is impaired during co-infection. (A) Volcano plots of differential gene expression in inner versus outer region myeloid cells in granulomas among groups. Each point represents one gene; horizontal dotted lines indicate adjusted p-values at 0.05 and vertical lines indicate 1 and -1 log-fold change. (B) Log2-transformed expression of genes involved in tryptophan catabolism among all granuloma myeloid cells. Genes from the GSEA leading edge. (C) Log2-transformed expression of genes involved in tryptophan catabolism between inner versus outer regions of granuloma myeloid cells. Genes from the GSEA leading edge, only adjusted p-values <0.05 are listed. (D) Log2-transformed expression of genes involved in reactive oxygen and nitrogen detoxification in inner versus outer regions of granuloma myeloid cells. Genes from the GSEA leading edge, only adjusted p-values <0.05 are listed. (E) GSEA (redundancy reduced) comparing myeloid cells in the inner region by group. Pathways are from WebGestalt using the Reactome database. Only pathways with adjusted p-values < 0.5 are shown.

One pathway seen in the Mtb inner ring of myeloid cells (Supplemental Table 7) and previously described in TB was the metabolism of the essential amino acid tryptophan (22, 23). Starvation of tryptophan in the local environment is one mechanism by which myeloid cells can kill intracellular pathogens (24). Indoleamine 2,3-dioxygenase 1 (IDO1) is the rate-limiting enzyme that mediates the breakdown of tryptophan. Overall IDO1 expression was upregulated in SIV/Mtb granulomas compared to both SIV/ART/Mtb and Mtb only granulomas (Figure 4B). When assessed spatially, genes from multiple steps in catabolism of tryptophan were upregulated specifically in myeloid cells in the inner ring of Mtb only granulomas and downregulated in the outer ring (Figure 4C), as previously observed (20). However, the spatially distinct differential expression of most genes was absent in myeloid cells between the inner and outer rings in both SIV/ART/Mtb and SIV/Mtb granulomas, which had similarly high expression of these genes throughout the granuloma and in which the outer ring lacked downregulated expression (Figure 4C). A similar pattern of expression was also seen for reactive oxygen/nitrogen species (ROS/RNS) detoxification, another mechanism by which myeloid cells can eliminate intracellular pathogens (24). Myeloid cells from the inner ROI had higher expression of genes in the ROS/RNS detoxification pathway compared to outer region ROIs of Mtb only granulomas (Figure 4D). In contrast, the distinction between inner and outer myeloid cell ROIs from SIV/Mtb and SIV/ART/Mtb granulomas in this pathway were lost (Figure 4D), suggesting that these granulomas do not restrict expression of these genes to the inner region as observed in Mtb only granulomas. Finally, expression patterns in myeloid cell ROIs from the inner and outer regions were compared between groups directly (Figure 4E, Supplemental Figure 5, Supplemental Tables 8-11). GSEA showed that DEGs in the myeloid cells in the inner region of SIV/Mtb granulomas, where the majority of myeloid cells are located, were enriched in collagen biosynthesis and pro-inflammatory pathways, including type I interferon signaling, complement responses and cytokine signaling, compared to both SIV/ART/Mtb and Mtb granulomas (Figure 4E, Supplemental Table 9). In contrast, Mtb only myeloid cells in the inner region were enriched in DEGs from pathways related to gene expression and protein synthesis (translation initiation and elongation) (Figure 4E, Supplemental Table 9). Thus, the overall increased pro-inflammatory environment in the inner region of granulomas during SIV co-infection is driven by myeloid cells within the inner region, which is not corrected by early suppressive ART.

A parallel analysis of the spatial pattern among the inner and outer rings of granuloma T cells in each group was performed. As with myeloid cells, T cells in the inner and outer regions of Mtb only granulomas had very distinct gene expression patterns from each other (981 differentially expressed genes; Figure 5A, Supplemental Table 12). T cells in the inner region were dominated by enrichment in DEGs from pathways related to metabolism and biosynthesis, whereas the outer ring was enriched in protein synthesis DEGs (Supplemental Table 13). This spatial pattern was impaired in SIV/Mtb and SIV/ART/Mtb granulomas, in which a more homogenous gene expression pattern was observed with fewer differentially expressed genes between the inner and outer rings (Figure 5A). Gene expression from T cells SIV/ART/Mtb granulomas were less spatially distinct than Mtb only granulomas, with 439 differentially expressed genes (Supplemental Table 12), though the pattern of DEG enrichment was similar: an inner region of metabolism and biosynthesis and outer ring of protein synthesis (Supplemental Table 13). Finally, only 9 differentially expressed genes were identified between the inner and outer rings of SIV/Mtb granuloma T cells, with similar expression in the inner and outer regions and with no differentially expressed pathways (Figure 5A, Supplemental Tables 12-13).

**Figure 5.**
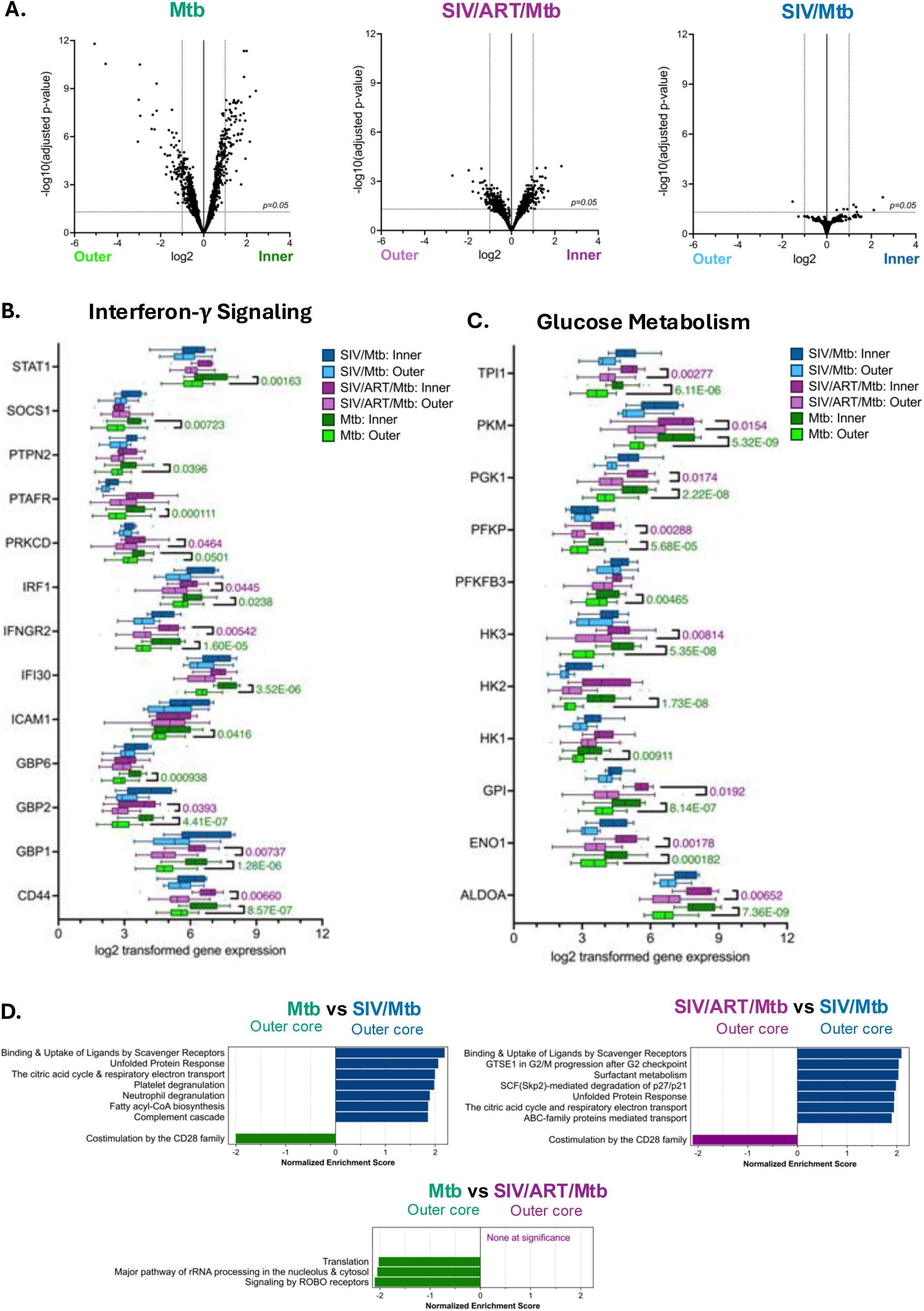
Spatial organization of T cell function in granulomas is impaired during co-infection. (A) Volcano plots of differential gene expression in inner versus outer region T cells in granulomas among groups. Each point represents one gene; horizontal dotted lines indicate adjusted p-values at 0.05 and vertical lines indicate 1 and -1 log-fold change. (B) Log2-transformed expression of genes involved in interferon-γ signaling between inner and outer region T cells in granulomas. Genes from the gene set enrichment analysis leading edge, only adjusted p-values <0.05 are listed. (C) Log2-transformed expression of genes involved in glucose metabolism between inner versus outer region T cells. Genes from the gene set enrichment analysis leading edge, only adjusted p-values <0.05 are listed. (D) GSEA (redundancy reduced) comparing T cells in the outer region by group. Pathways are from WebGestalt using the Reactome database. Only pathways with adjusted p-values < 0.5 are shown.

This spatial distinction was exemplified in the interferon-γ signaling pathway, among the most enriched immune DEGs in inner ring T cells, in which T cells in the inner region had higher expression of interferon-γ signaling genes than the outer ring in Mtb only granulomas (Figure 5B). Some, but not all, T cell genes in the interferon-γ pathway in SIV/ART/Mtb granulomas had a similar differential pattern as Mtb only T cells (Figure 5B). However, this differential pattern of expression was not observed in SIV/Mtb granuloma T cells, where gene expression was similar in the inner and outer rings (Figure 5B). Upstream of this interferon-γ signaling, there was an overall overexpression of most genes related to interferon-γ production in T cells in Mtb only granulomas relative to SIV/Mtb, though this pattern was not spatially distinct (Supplemental Figure 6). The spatial pattern of glucose metabolism was similarly differentiated in Mtb only T cells, with higher expression in the inner ROIs than the outer ROIs, whereas the SIV/Mtb group had similar gene expression levels in both regions (Figure 5C). Lastly, the SIV/ART/Mtb also had an intermediate phenotype where expression of some genes in the glucose metabolism pathway was differentially expressed by region and others were not (Figure 5C).

In addition to comparing gene expression of T cells the inner versus outer regions in each group, the gene expression patterns of T cells in the outer and inner granuloma regions were compared directly between groups (Figure 5D, Supplemental Figure 7, Supplemental Tables 14-17). In the outer ring of the granuloma, where the majority of T cells are localized, SIV/Mtb T cells were enriched in DEGs from metabolic (respiratory electron transport, fatty acyl-CoA biosynthesis) and immune pathways (complement cascade, type I interferon) (Figure 5D). Whereas T cells from the outer ring of Mtb only and SIV/ART/Mtb granulomas were enriched in DEGS from pathways related to T cell activation and signaling (CD28 co-stimulation) compared to SIV/Mtb outer T cells ROIs (Figure 5D). This decrease in T cell signaling and activation in SIV/Mtb granulomas driven by the outer region T cells may be associated with perturbed interactions with infected macrophages in granulomas, and thus contributes to increased bacterial burden compared to granulomas in animals infected with only Mtb.

### SIV RNA within co-infected granulomas is present within both macrophages and T cells

SIV can infect cells with granulomas (11, 12), and both SIV and HIV can alter the function of both macrophages and T cells (25–32). However, the extent to which SIV infects macrophages versus T cells in granulomas is unclear. To assess which cells were SIV-infected and where in these granulomas SIV-infected cells were localized, serial sections of granulomas were stained with RNAscope probes specific for SIV as well as markers for macrophages (CD68 and CD11c) and T cells (CD3). SIV-infected cells were identified as triple positive for SIV RNA, DAPI, and their respective cell markers. SIV/Mtb granulomas had higher viral RNA frequencies, which were similar in macrophages and T cells, with a median of 44% of macrophages and 38% of T cells containing detectable SIV RNA with some heterogeneity among individual granulomas (Figure 6, Supplemental Figure 8). SIV/ART/Mtb granulomas, in which SIV RNA was undetectable by qRT-PCR (16), had lower detectable SIV RNA frequencies in both macrophages (median 3%) and T cells (median 5%) (Figure 6, Supplemental Figure 8). SIV RNA was also throughout the granuloma, from the outer lymphocyte cuff into the myeloid region (Figure 6A). As SIV can infect both macrophages and T cells in granulomas, the function of both cell types may be perturbed by viral infection, which is likely associated with the differences in both myeloid and T cell transcriptional changes throughout the granuloma and poor control of Mtb burden in granulomas.

**Figure 6.**
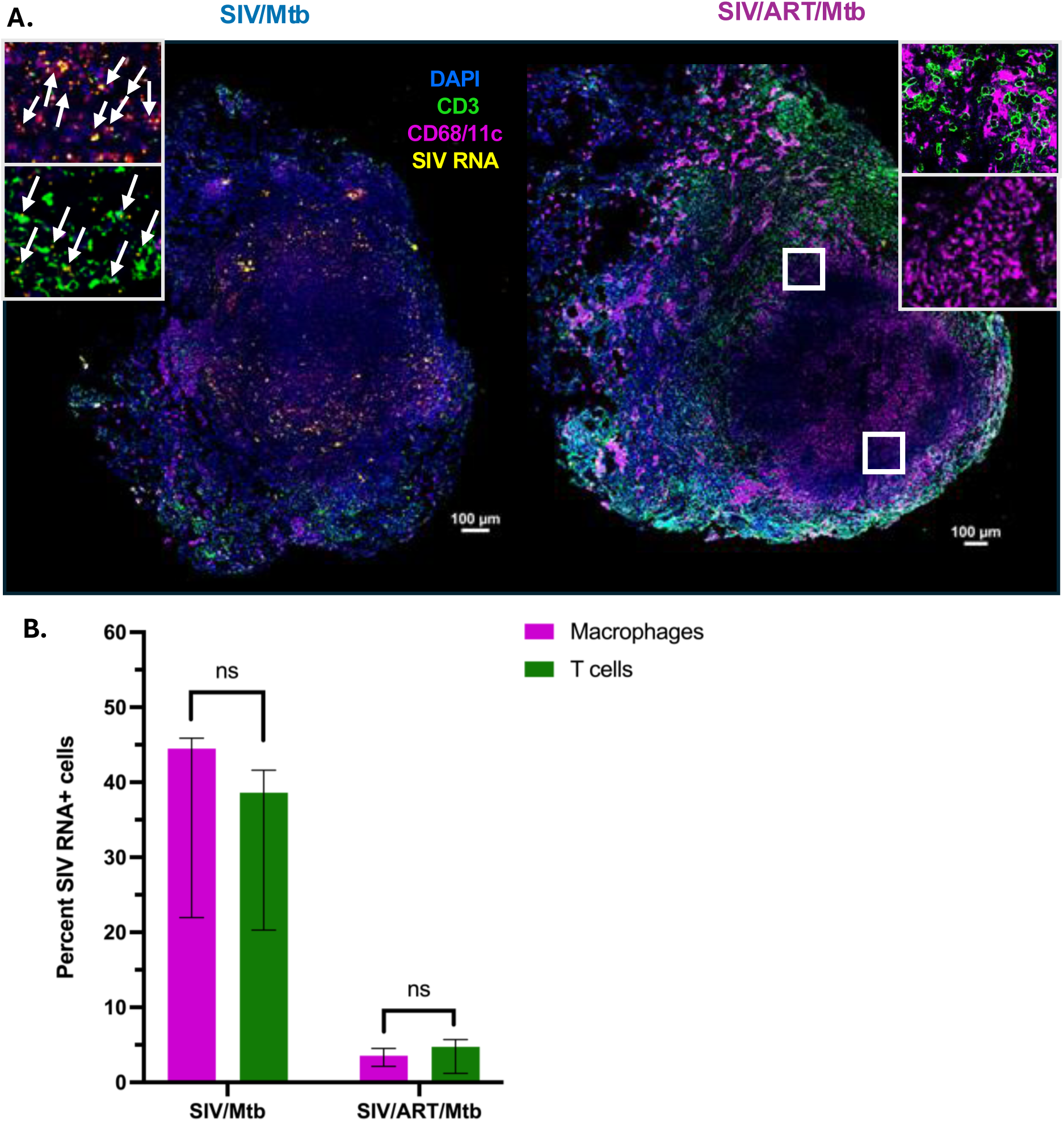
SIV infection within co-infected granulomas is distributed in both myeloid cells and T cells. (A) RNAscope (SIV *gag* and *pol*; yellow) and immunohistochemistry (CD3+ T cells, green; CD11c/CD68+ myeloid cells, magenta) staining in SIV/Mtb (left) and SIV/ART/Mtb (right) granulomas. (B) Median percent with interquartile range of the percent myeloid cells (CD68+CD11c+DAPI+; magenta bars) and the percent of T cells (CD3+DAPI+; green bars) that had detectable SIV RNA. N= 7 granulomas from 5 SIV/Mtb animals (1-2 granulomas per animal) and 7 granulomas from 6 SIV/ART/Mtb animals (1-2 granulomas per animal).

### Spatial analysis of ligand-receptor pair communication within granulomas

To understand how macrophages and T cells interact with each other and how that communication could be altered by SIV in granulomas, cell-cell communication was assessed using CellChat. CellChat compares gene expression of ligands and receptors in cells and uses a database of known ligand-receptor pairs to determine which cells are likely to be communicating with each other (33). ROIs were grouped into inner macrophages, outer macrophages, inner T cells, and outer T cells to compare communication in granulomas across groups in a spatial context. The cellular communication network of these ROIs was visualized by mapping the strength of ligand-receptor interactions between cell types, with interaction strength defined by the degree of expression of ligands and receptors. This spatially informed communication network showed that the network edges, which represent the strength of interactions between ROIs, were weaker in SIV/Mtb granulomas, particularly between inner T cells and both inner and outer macrophages, compared to the strength of interactions between inner and outer macrophages, which was comparable between groups (Figure 7A-C). Communication strength in SIV/ART/Mtb granulomas had an intermediate phenotype, with stronger communication between inner T cell ROIs and inner and outer macrophage ROIs than in SIV/Mtb granulomas but not as strong as Mtb only granulomas (Figure 7A-C). This pattern in the reduction in strength of cell-cell communication in the context of SIV/Mtb co-infection was also true when considering the number of interactions between cell groups, particularly inner region T cells communication with outer and inner region macrophages (Supplemental Figure 9A-C).

**Figure 7.**
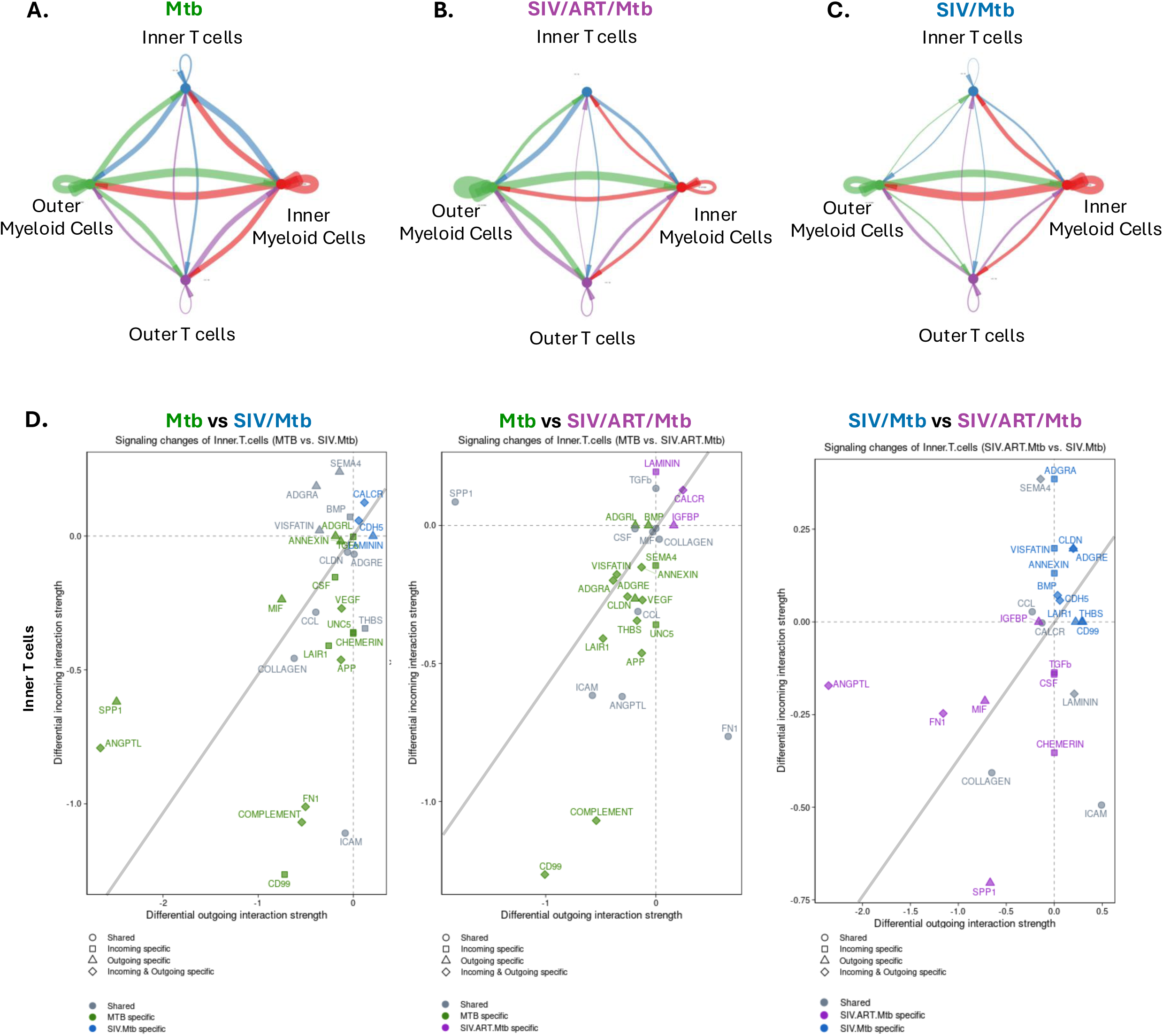
Analysis of cell-cell communication in granulomas. (A-C) Aggregated cell-cell communication networks between inner and outer myeloid cells and T cells in (A) Mtb only granulomas, (B) SIV/ART/Mtb granulomas, and (C) SIV/Mtb groups. Arrows indicate signal senders to signal receivers. Colors indicate sender group: green is outer myeloid cells, red is inner myeloid cells, purple is outer T cells, and blue is inner T cells. Arrow thickness is proportional to the strength of ligand-receptor interactions. Node size is proportional to the number of ROIs in that cell type. (D) Ligand-receptor signaling strength changes in inner T cells summarized as pathways in each paired comparison of Mtb vs. SIV/Mtb (left), Mtb vs. SIV/ART/Mtb (middle), and SIV/Mtb vs. SIV/ART/MTB (right) groups. Each symbol indicates a pathway, color indicates group, shape indicates signal direction.

To further characterize these defects in cellular communication and their directionality, the summative incoming and outgoing interaction strength of sending and receiving spatially defined cell types were visualized. While T cells throughout the granuloma had lower overall interactions relative to inner and outer macrophages, both incoming and outgoing signaling in T cells in the inner region specifically were decreased in the SIV/Mtb and SIV/ART/Mtb granulomas compared to Mtb only granulomas (Supplemental Figure 9B). Inner T cells are in closer proximity to the Mtb-infected macrophages in the inner region, and thus, reduction in their signaling to macrophages may reflect a mechanism of impaired bacterial control during SIV/Mtb co-infection. To identify the signaling pathways that comprised this defect in communication from the inner region T cells, pathways with differential ligand-receptor signaling in each spatially informed cell type were compared across groups (Figure 7D). Inner region T cells in Mtb only granulomas were enriched in DEGs from signaling pathways related to cellular migration, adhesion, and migration relative to SIV/Mtb inner region T cells, including signaling through intracellular adhesion molecule (ICAM), fibronectin (FN1), adhesion G-protein couple receptors (ADGRL), and thrombospondin (THBS) (Figure 7D). Signaling in these pathways was also reduced to a lesser extent compared to SIV/ART/Mtb inner region T cells (Figure 7D). The decreased communication in these SIV/Mtb granulomas between inner T cells and both inner and outer myeloid cells as well as among T cells is consistent with GSEA data demonstrating a reduction in T cell signaling pathways. Decreased signaling in these pathways may also reflect dysregulation in myeloid cell and T cell interactions, and thus a greater opportunity for bacterial dissemination and increased bacterial burden, which may be correlated with the increase in extrapulmonary disease seen in SIV/Mtb and SIV/ART/Mtb animals (16).

## Discussion

The granuloma is the centerpiece of immune cell interactions during Mtb infection, in which immune cells are recruited to the site of infection in a highly organized architectural structure, starting from a central area of Mtb-infected myeloid cells, such as macrophages and dendritic cells, with or without necrosis surrounded by peripheral zones of lymphocytes. Insights into the complex interactions within the highly organized immune landscape of the granuloma (both regional and cell-to-cell) have only recently been undertaken with the advent of novel spatial, systems immunology and computational techniques. In this study, IDO1 expression was limited to the inner myeloid region of Mtb granulomas. Similar to our findings, granulomas from Mtb infected NHPs and humans have been shown to produce IDO1 within the central myeloid areas (20, 22, 34), and closely positioned near the lymphocytic cuff. Some have speculated that IDO1 may play a key role in immune suppression near the hypoxic central area of the granuloma, an area of high Mtb burden, to prevent excessive inflammation (19). This area of hypoxia may also be a dominant driver of the complex immune interactions at this interface between the inner and outer rings of granulomas observed in our study. Our work contributes to the field by highlighting the functionally distinct nature of each zone composing the granuloma during Mtb infection and demonstrating that this distinction is lost during SIV-Mtb co-infection that is associated with high Mtb burden and poor clinical outcome.

The HIV/Mtb syndemic continues to be major cause of morbidity and mortality worldwide (1, 3), and yet understanding the mechanisms that lead to worse outcomes in co-infection are difficult to study in humans, as infection order and duration is typically unknown or imprecise and relevant tissue cannot easily be obtained antemortem. We used an NHP model of HIV/Mtb co-infection that recapitulates human infection where Mtb and SIV infection as well as ART administration are highly controlled. We and others have shown that SIV RNA copies can be found in granulomas (12, 35), and our data further show that SIV infects both myeloid cells and T cells within granulomas at similar frequencies. Our data instead emphasizes that co-infection leads to a disruption in the zonal organization of cell function and cell-cell communication rather than a disruption of granuloma architecture. SIV/Mtb granulomas exhibit more pro-inflammatory environments, with enrichment in DEGs from pathways involving type 1 interferon signaling, than Mtb only or SIV/ART/Mtb granulomas, particularly within myeloid cells. T cells in SIV/Mtb granulomas were less enriched in DEGs from activation and downstream signaling pathways, and had an overall reduction in communication among myeloid cells and T cells, particularly in signaling pathways related to collagen, cell migration, and cell adhesion, which were only partially restored in SIV/ART/Mtb granulomas, which lack viral replication.

Type I interferon (interferon α and β) signaling is a canonical antiviral immune response that mediates the upregulation of interferon-stimulated genes (ISGs) through JAK/STAT signaling (36). These responses are upregulated during HIV infection (36, 37) and are necessary to stimulate innate immune responses early in infection (38). However, Type I interferon signaling in TB is less understood and has been correlated with more severe outcomes of infection and higher bacterial burden (39–42). Moreover, sustained type I interferon leads to a compensatory increase in suppressive IL-10 and dysfunctional T cells, impairing the balance necessary for successful clearance of Mtb (43). In the present study, gene expression from type I interferon signaling was enriched in myeloid cells in SIV/Mtb granulomas relative to both Mtb only and SIV/ART/Mtb granulomas, suggesting that unsuppressed viral replication is associated with a more pronounced pro-inflammatory response. Whether this causes higher Mtb burden or results from that higher Mtb burden is unclear from this single timepoint study. A pro-inflammatory milieu may skew the phenotype of activated macrophages, which are polarized into M1 or M2 phenotypes based on the local microenvironment, such that M1 macrophages are pro-inflammatory while M2 macrophages are anti-inflammatory and associated with wound healing (44). Studies using a combination of NHP samples and in silico modeling have demonstrated that an initial pro-inflammatory environment in granuloma macrophages is necessary for eventual bacterial containment of Mtb, but that prolonged pro-inflammatory polarization is detrimental for controlling dissemination (45). Thus, sustained pro-inflammatory interferon signaling in the SIV/Mtb myeloid cells compared to the Mtb only and SIV/ART/Mtb myeloid cells may contribute to their higher bacterial burden, though further studies are necessary to determine causality.

Tryptophan catabolism has been recognized to play an important role in TB granulomas and spatially organized tryptophan catabolism in Mtb granulomas has also been observed previously in granulomas during Mtb-only infection (20, 22). Tryptophan catabolism is immunosuppressive: it skews T cell polarization towards regulatory T cells and reduces T cell proliferation (46, 47), and can promote M2 polarization in myeloid cells (48). IDO1, the rate limiting enzyme in tryptophan catabolism, has also been studied as a potential biomarker for TB disease in humans (49, 50). While IDO1 inhibition reduced bacterial burden during early Mtb infection in a highly susceptible macaque model (22), it did not alter burden during SIV/Mtb co-infection during ART despite improved quality of immune responses (21). In this study, SIV/Mtb granulomas did not have downregulated IDO1 expression in the outer region and instead had an altered spatial pattern with higher levels of IDO1 throughout the granuloma. The role of IDO1 in Mtb outcomes is likely multifactorial and complex as it has been observed in granulomas with both non-viable and viable Mtb (19).T cell-derived interferon-γ, also known as type 2 interferon, has been the focus of T cell-mediated immunity to TB [reviewed in (51)]. CD4+ T cell secretion of interferon-γ activates macrophages for elimination of phagocytosed pathogens [reviewed in (52)], and has been shown to be important in the control of Mtb infection in mice (53, 54) though interferon-independent mechanisms of control have also been demonstrated (55, 56). TB vaccine studies using peripheral interferon-γ production as a correlate of protection have not yielded consistent results (57), though recent studies in NHP have shown that airway T cell production of interferon-γ may be a better predictor of protective? immunity (58). In this study, expression of genes in the interferon-γ signaling pathway, but not interferon-γ production pathway, in granuloma T cells was spatially differentiated within the inner and outer regions of Mtb only granulomas but not spatially differentiated within SIV/Mtb granulomas. This added context provides insight into how the spatial distribution of effector interferon-γ signaling may play a larger role in infection outcomes. Also, glucose metabolism in T cells was spatially differentiated in Mtb only but not SIV/Mtb granulomas. Glucose metabolism by T cells has been linked to their activation and effector function [(59) and reviewed in (60)]. These findings are also consistent with differences in glucose metabolism observed between Mtb only and SIV/Mtb groups, as T cell activation is associated with shifts in glucose metabolism (the “Warburg effect”) as oxidative phosphorylation and fatty oxidation pathways shift to aerobic glycolysis despite available oxygen [reviewed in (61)]. Infection of CD4+ T cells by HIV has been shown to increase the metabolism of glucose in T cells (62), and HIV replication is enhanced by cellular use of glycolysis (63). In the co-infected granulomas in this study, SIV can be found throughout both myeloid and T cell regions of the granuloma, which may be related to the lack of spatial restriction of enrichment of glucose metabolism pathway genes to the outer ring in SIV/Mtb granulomas. In both these T cell pathways, ART did not restore the spatial distinction of all gene expression. The data from this study highlight how granuloma success or failure in bacterial containment is associated with immune cell interactions in the granuloma, particularly between myeloid cells and T cells, and provides better understanding of the spatial context of that communication. Other studies investigating the spatial dynamics of Mtb granulomas have also shown that granulomas are highly organized, and that protective responses to Mtb in granulomas are not a dichotomous beneficial anti-inflammatory or harmful pro-inflammatory environment. Instead, studies have shown that an initial pro-inflammatory response is critical to bacterial control, but that sustained pro-inflammatory responses are detrimental to granuloma control of Mtb (45, 64). This study demonstrates how SIV co-infection impairs the organization of cellular function within the granuloma, such that SIV/Mtb granulomas are overall more inflammatory environments throughout the entire granuloma and antimycobacterial functions are no longer contained to the inner region as in Mtb only granulomas. This organization may be required to prevent ongoing immunopathology as has been seen in other Mtb granulomas (9).

We also show that ART can restore some but not all aspects of cellular functional impairment in granulomas. ART-treated animals did not have granulomas with spatially differentiated cellular function to the same extent as Mtb granulomas, but they had less inflammatory environments, stronger cell-cell communication and ultimately lower bacterial burden relative to granulomas from ART-naive SIV/Mtb animals. This suggests that SIV infection in the absence of ongoing replication with very early ART suppression still alters the spatially distinct regions of the granuloma. By demonstrating that SIV infects macrophages as well as T cells within granulomas, this study also provides another potential source of cellular dysregulation that may be associated with poor bacterial control.

This study was limited in that the GeoMX platform is not a single-cell assay, and as such the resolution per ROI was a median of 187 cells. Therefore, the conclusions should be interpreted as changes within groups of cells in granuloma regions rather than changes in individual cells. Cell-cell communication was assessed by labeling ROIs as in either the inner or outer ring, which limits resolution in spatial analysis. Due to the limitations of the number of antibodies for immunophenotyping with this platform, CD68 was used to define all myeloid cells. Myeloid cell ROIs in this study also likely include nearby neutrophils in addition to macrophages. The differential expression of S100A8 and S100A9 proteins, classically though not exclusively found in neutrophils (65), further supports that this myeloid cell subset includes neutrophils as well as macrophages. Thus, these findings should be considered to reflect changes in both cell types. Finally, our data includes all CD3+ cells in the T cell subset and does not distinguish between CD4+ and CD8+ T cells. The frequency of CD4+ T cells was significantly reduced in the granulomas in this study, which was not observed in SIV/ART/Mtb granulomas (16). As expected, the frequency of CD8+ T cells increased relative to CD4+ T cells in the SIV/Mtb group (16), which could confound the T cell profiles. The SIV/Mtb group in this study contained a fewer number of T cell ROIs, especially in the outer region, as expected, though all T cell ROIs from all groups were filtered during quality control such that they contained equivalent numbers of T cell nuclei (Supplemental Table 18). Thus, the reduction in global T cell signaling cannot be attributed to a specific T cell subset but rather to T cells overall. Lastly, while we can identify SIV+ cells in granulomas, identifying viable Mtb on a single cell basis is not yet technologically available.

This study raises important questions about the cellular interactions in the granuloma that may be important for controlling TB disease burden in the context of HIV co-infection, including during suppressive ART. Newer spatial transcriptomics platforms that can more closely approximate single-cell resolution will be useful to disentangle the interactions between specific subsets of myeloid and T cells. Further work is needed to understand whether modulation of differentially expressed pathways in this analysis can rescue cell-cell communication in the context of viral co-infection and may lead to development of targets for host-directed therapies and vaccine development.

## Materials and Methods

### Animal care and infections

Chinese cynomolgus macaques aged 5-8 years (Supplemental Table 1) were purchased from Valley Biosystems (Sacramento, CA) and randomized to Mtb-only (N=10), SIV/Mtb (N=9), or SIV/ART/Mtb (N=10) infection groups. SIV-infected macaques were infected with 7x10^4^ infectious units (IU) SIV_ΔB670_ (ARP-633, HIV Reagent Program) intravenously and randomized to either daily ART (tenofovir [20 mg/kg], emtricitabine [40 mg/kg], and dolutegravir [2.5 mg/kg] subcutaneously) beginning three days post-SIV infection or no ART. SIV/Mtb animals (at 4 months of SIV infection) and Mtb-only animals were infected with low-dose (8-17 CFU) barcoded Mtb Erdman strain (66) via bronchoscopic instillation as previously described (67, 68). Animals were housed at a BSL-3 NHP facility in the Regional Biocontainment Laboratory at the University of Pittsburgh after Mtb infection. Mtb infection was confirmed by presence of granulomas on serial PET-CT as previously described (69–71). SIV and Mtb disease progression was monitored by serial peripheral blood draw (plasma viremia, immune responses, erythrocyte sedimentation rate), bronchoalveolar lavage (immune responses, Mtb growth), gastric aspirate (Mtb growth), PET-CT, and daily clinical examination.

### Sex as a biological variable

This study encompassed samples from 21 male NHPs and 2 female NHPs (Supplemental Table 1) due to limitations in availability of female NHPs.

### Necropsy, tissue processing, Mtb quantitation, and SIV quantitation

Animals were necropsied as previously described (72). In summary, PET-CT matched granulomas from the lung, lymph node, and other sites as well as grossly appearing lung and lymph node, were excised and assessed for immune responses, histopathology, bacterial burden, and viral burden. TB lesions and other tissues were divided in half such that was placed in formalin after excision then processed into formalin-fixed paraffin-embedded (FFPE) blocks and half was homogenized for assessment of bacterial burden, viral burden, and immune responses. To quantify Mtb burden, tissue homogenates were plated on 7H11 plates to assess colony-forming units (CFU) and estimated CFU per granuloma was calculated as previously described (72). Homogenates were also stored in Trizol LS for assessment of viral burden, and then RNA was extracted as previously described (16). SIV in tissue was quantified by qRT-PCR also as previously described [(16, 73, 74)] using SuperScript III First-Strand Synthesis SuperMix (Thermo Fisher) for cDNA synthesis, SsoAdvanced Universal Probes Supermix (BioRad) for qPCR, with primers and probes for SIV_ΔB670_ (forward 5’-GTCTGCGTCATTTGGTGCATTC-3’, reverse 5’-CACTAGATGTCTCTGCACTATTTGTTTTG-3’, probe FAM-CGCAGAAGAGAAAGTGAAACATACTGAGGAAG-TAMRA). Tissue SIV RNA was normalized per 10^6^ CD4 RNA. CD4+ T cell frequencies in granulomas were measured with flow cytometry as previously described (16). Briefly, granuloma single cell homogenates were incubated with media and Brefeldin A for four hours and stained with antibodies as previously described (16). Samples were run on an LSR II (BD) and analyzed with FlowJo Software v.10.8 (Treestar Inc, Ashland, OR).

### Nanostring GeoMX assay

Freshly cut tissue sections 5 mM thick were cut from lung granuloma FFPE blocks (Supplemental Table 1) less than three years old (SIV/Mtb N = 7, SIV/ART/Mtb = 8, Mtb = 9 granulomas) and mounted onto positively charged slides (Fisherbrand Superfrost Plus slides). Granulomas were selected for which there was associated metadata (CFU, SIV RNA, histology; Supplemental Table 1) and which came from different animals within each group. Slides were stained with CD68 (Thermo Fisher, mouse clone KP1), CDE3 (Origene, mouse clone UMAB54), CD8a (Origine, mouse clone OTI3H6), and the Nanostring GeoMX Human Whole Transcriptome Atlas barcoded RNA probes. Probes bind to complimentary host mRNA in tissue, and then the probes are selectively UV-photocleaved for collection and sequencing using masks based on antibody cell type markers to assess the transcriptome from specific cell types in the specific ROIs in a tissue (75). There is high homology between the macaque and human genomes (76), and granuloma ROI samples were well above the minimum suggested raw read and aligned read counts, supporting high confidence in cross-reactivity between NHP and human RNA. Regions of interest (ROIs) in granulomas were chosen based on antibody staining as well as hematoxylin & eosin (H&E) stained serial sections. RNA probes from cell-type specific ROIs were photocleaved were sequenced using Nanostring’s nCounter and Digital Spatial Profiler platforms. Raw data were analyzed and quality checked using the NanoStringNCTools, GeomxTools (version 3.10.0), and GeoMxWorkflows (version 1.0.1) packages available via Bioconductor (version 3.20) using R (version 4.22) and RStudio (version 2023.12.1+402). All ROIs were filtered to include a minimum of 1000 reads, 80% trimmed reads, 80% stitched reads, 80% aligned reads, 50% sequencing saturation, and 50 µm^2^ area. Myeloid cell ROIs were additionally filtered to contain between 67-670 nuclei, and lymphocyte ROIs were filtered to contain between 70-797 nuclei (maximum ten-fold range). ROIs with expressing less than 5% of genes above the limit of detection and genes detected in fewer than 20% of ROIs were filtered out.

### Batch correction, differential gene expression, and pathway analyses

Cell-type specific filtered GeoMx data were batch corrected and differential gene expression was assessed using the DESeq2 (version 1.28.3) package available via Bioconductor using R (version 4.22) and RStudio (version 2023.12.1+40 2). Data generated from two GeoMx runs were batch corrected in DESeq2 by including batch as the first factor in the analysis design, log2 transforming using variance stabilization, and using removeBatchEffect function in limma. Differential gene expression per cell type in granulomas overall was determined using the Wald statistical test where the design was ∼ batch + treatmentgroup. Spatially defined differential gene expression per cell type (i.e., inner ROIs versus outer ROIs) within each group was determined using the Wald test where the design was ∼ batch + location. Spatially defined differential gene expression per cell type within each region (i.e., inner ROIs compared across groups) among each group was determined using the Wald test where the design was ∼ batch + treatmentgroup. Statistical significance for all Wald tests was set at an adjusted p-value < 0.05 Pathway analysis was assessed using GSEA was performed using WEB-based Gene SeT AnaLysis Toolkit (WebGestalt) (version 2019) using Reactome and Gene Ontology databases with statistical significance set at a false discover rate < 0.05. Differentially expressed genes from the leading edge of the pathways were used for visualization.

### Cell-cell communication analysis

Cell-cell communication in lung granulomas was assessed using the CellChat package, (version 2.1.2, https://github.com/jinworks/CellChat) using R (version 4.4.0) and RStudio (version 0.5.0) via the University of Pittsburgh Center for Research Computing’s htc cluster to compare expression of ligands and receptors in ROIs. The batch-corrected log_2_-normalized gene expression matrix (gene x ROI) of genes with log_2_ normalized expression values above 0 from GeoMX was used as input. Macrophage and T cell ROIs were tagged in metadata as inner or outer ROIs based on hematoxylin & eosin images and GeoMX-stained immunohistochemistry images. In brief, communication probability for ligand-receptor pairs was assessed within in group using 5% truncated mean (i.e., the percent of ROIs in a group expressing a given ligand or receptor must be above 5%) and then combined into a merged CellChat object for data visualization per developer’s instructions. Overexpression of a signaling genes was determined using Wilcoxon rank sum test with significance set at an adjusted p-value < 0.05. Communication probability was determined as described in reference (33).

### RNAscope and immunohistochemistry

The presence of SIV_ΔB670_ RNA infected cells was visualized in granulomas using RNAscope® 2.5 HD Red (ACDbio). Tissue sections (5 uM thick) were cut from formalin-fixed paraffin embedded (FFPE) blocks and mounted onto positively charged slides (Fisherbrand Superfrost Plus slides). RNAscope® 2.5 HD Red chromogenic kit (Advanced Cell Diagnostics) was used to visualize SIV_ΔB670_ *gag* and *pol* mRNA as per the manufacturer’s instructions using ACDbio proprietary custom probes designed for SIV_ΔB670_ *gag* and *pol*. Briefly, slides were baked for 30 minutes at 60C before use and then deparaffinized in a xylene and ethanol immersion series. Slides were pretreated with hydrogen peroxide and ACDbio RNAscope® Protease Plus.

Microwave-based antigen retrieval was performed using tri-sodium citrate dihydrate buffer. Probes were hybridized to tissue and serially amplified as per the manufacturer’s instructions until signal detection. After signal detection using FastRed, tissues were subsequently stained with CD3 (Dako, rabbit polyclonal, 1:50), CD68 (Thermo Fisher, mouse clone KP1, 1:40), and CD11c (Leica, mouse clone 5D11, 1:40) overnight at 4°C. Secondary antibodies Alexa Fluor (AF) preabsorbed anti-rabbit 488, and AF preabsorbed anti-mouse 647 were used were used at 1:1000 dilution. Slide autofluorescence was quenched using Vector Labs TrueVIEW quenching kit per the manufacturer’s instructions (3 minute incubation post-secondary staining). Nuclei were visualized using DAPI in the mounting medium. Stitched images of granulomas were acquired using a widefield Nikon DS-Qi2 microscope (20X magnification, 0.75 aperture) and analyzed using the Nikon Elements High Content Analysis research package (version 6.10.01). Images were auto-deconvoluted and quantified using a General Analysis 3 algorithm to count DAPI+ cells, DAPI+CD3+ T cells, DAPI+CD68+CD11c+ macrophages, DAPI+CD3+SIV RNA+ infected T cells, and DAPI+CD68+CD11c+SIV RNA+ infected macrophages.

### Statistics

Statistical comparison of CD4+ T cells in lung granulomas and CFU in lung granulomas was assessed with a random effect model with the animal as the random effect and group and fixed effect), fit using the restricted maximum likelihood (REML) method (JMP Pro 17.2.0) and with Tukey HSD adjusted p-values. Statistical analyses for differential gene expression (DeSeq2), pathway analyses (GESA), and cell-cell communication probability (CellChat) were as described above. In all visualizations, only statistically significant (adjusted p-values < 0.05) comparisons are shown.

### Study approval

All animal protocols were approved by the University of Pittsburgh Institutional Animal Care and Use Committee in compliance with the Animal Welfare Act and Guide for the Care and Use of the Laboratory Animals.

## Data availability

The granuloma spatial transcriptomics data generated with Nanostring GeoMX and used in these experiments have been made publicly available through the Gene Expression Omnibus at GSE327094 (https://www.ncbi.nlm.nih.gov/geo/query/acc.cgi?acc=GSE327094).

## Author contributions

Z.A. and P.L.L. designed the research study, analyzed data, and wrote the manuscript. J.M.M. conducted experiments, acquired data, analyzed data, and wrote the manuscript. J.D. supervised data analysis and wrote the manuscript. J.M. provided reagents and analyzed data. P. M. analyzed data and conducted statistical analyses. P.S., C. K., C.R.D, T. R., E.K. conducted experiments, acquired data, and analyzed data.

## Acknowledgements

We thank all the excellent veterinary and laboratory staff at the Tuberculosis Research Group for their hard work and dedication to animal care and sample processing. This study would be impossible without their work. We thank E. Mike Meyer and the Nanostring core at the University of Pittsburgh Hillman Cancer Center for staining and sequencing the GeoMX samples and also thank Hanxi Xiao and Mary Cundiff from the Das lab for helpful conversations about systems immunology and bioinformatics analyses.

The authors have declared that no conflict of interest exists.

---

**Supplemental Figure 1.**
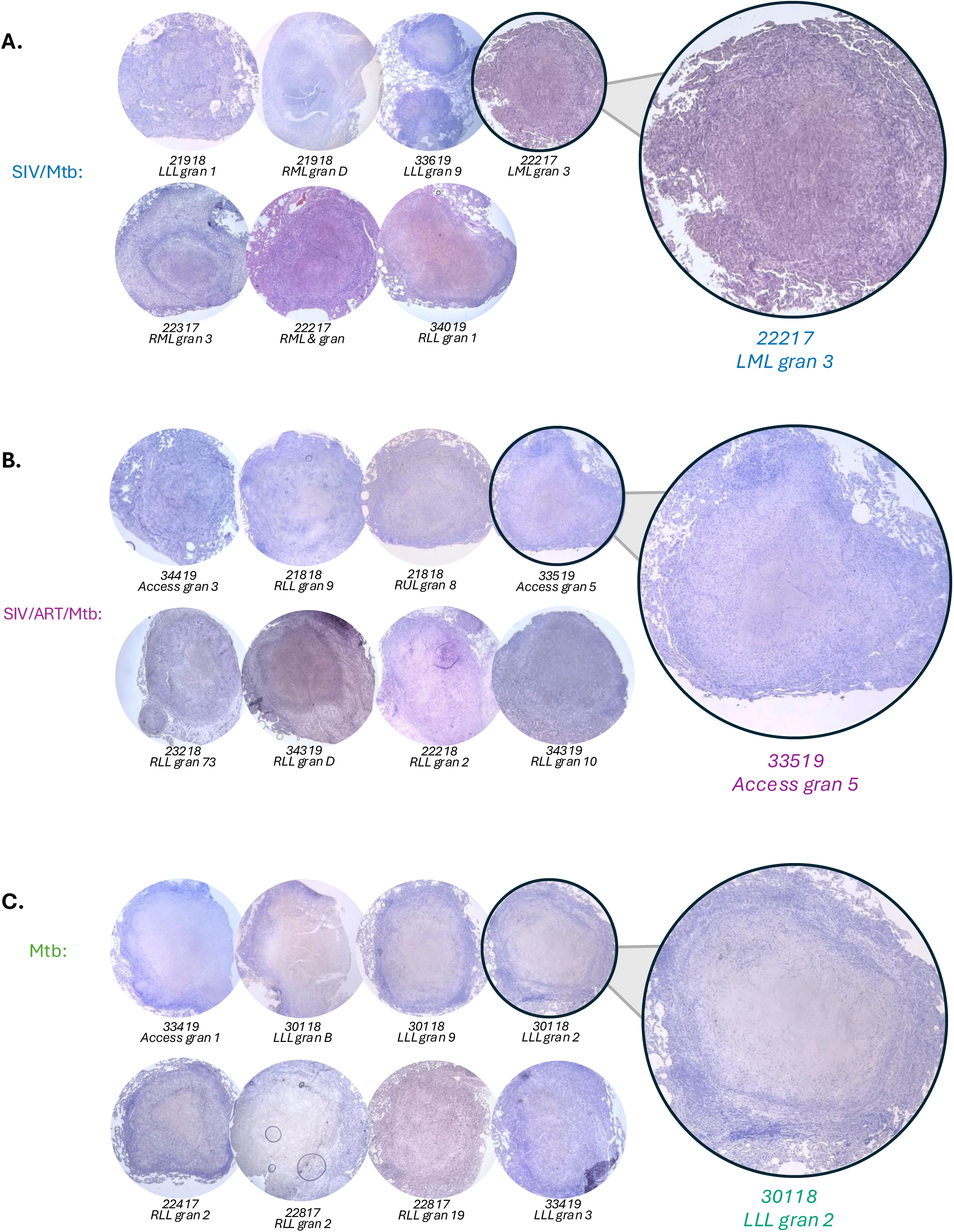
Granuloma histology does not differ between animal groups. Hematoxylin and eosin stained tissue lung granulomas (10X magnification) from SIV/Mtb (A), SIV/ART/Mtb (B), and Mtb (C) granulomas used in this study

**Supplemental Figure 2.**
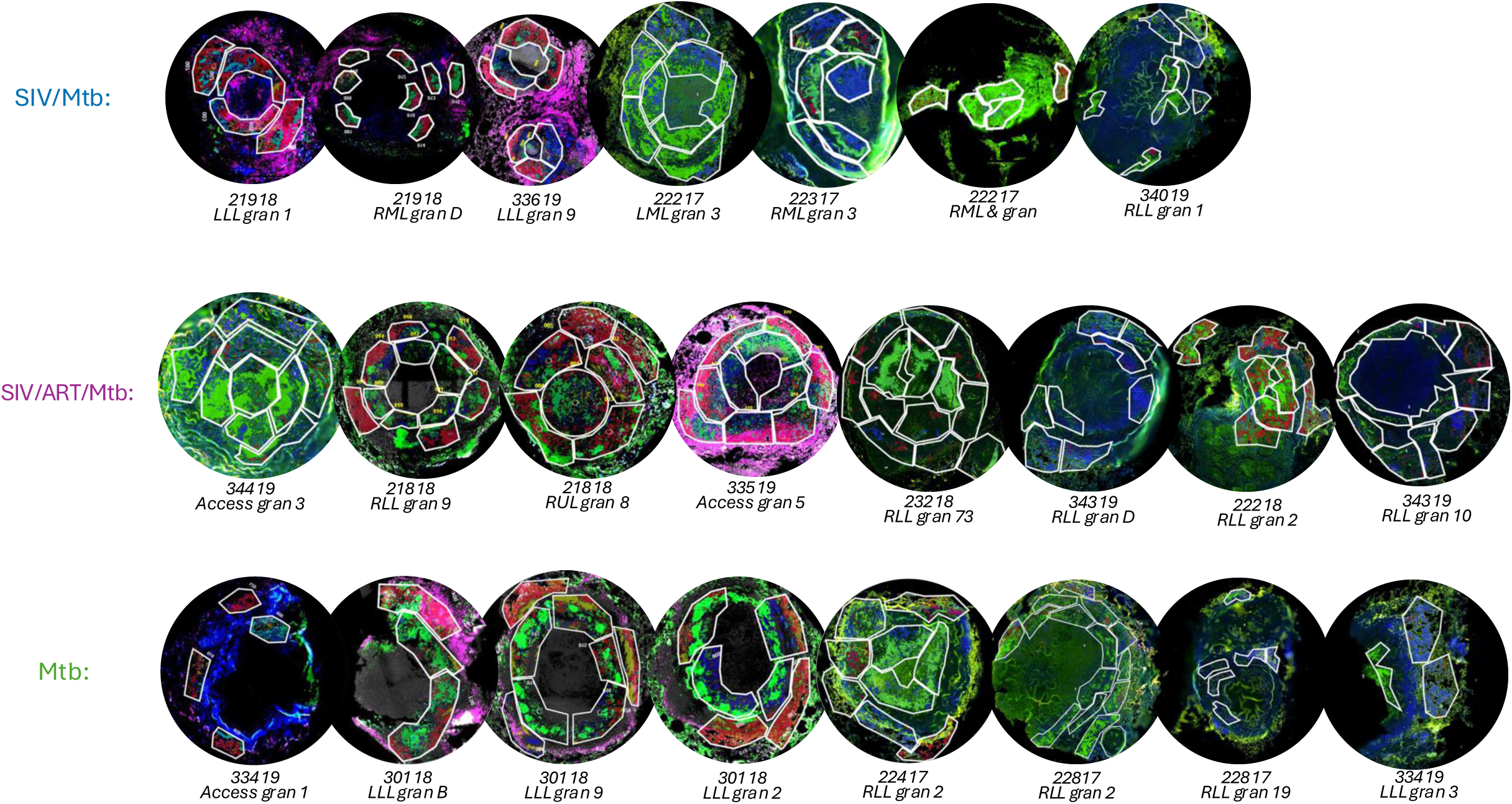
GeoMX regions of interest (ROIs) from granulomas. ROIs (inner and outer rings) from which spatial transcriptomics were assessed in SIV/Mtb (A), SIV/ART/Mtb (B), and Mtb (C) granulomas.

**Supplemental Figure 3.**
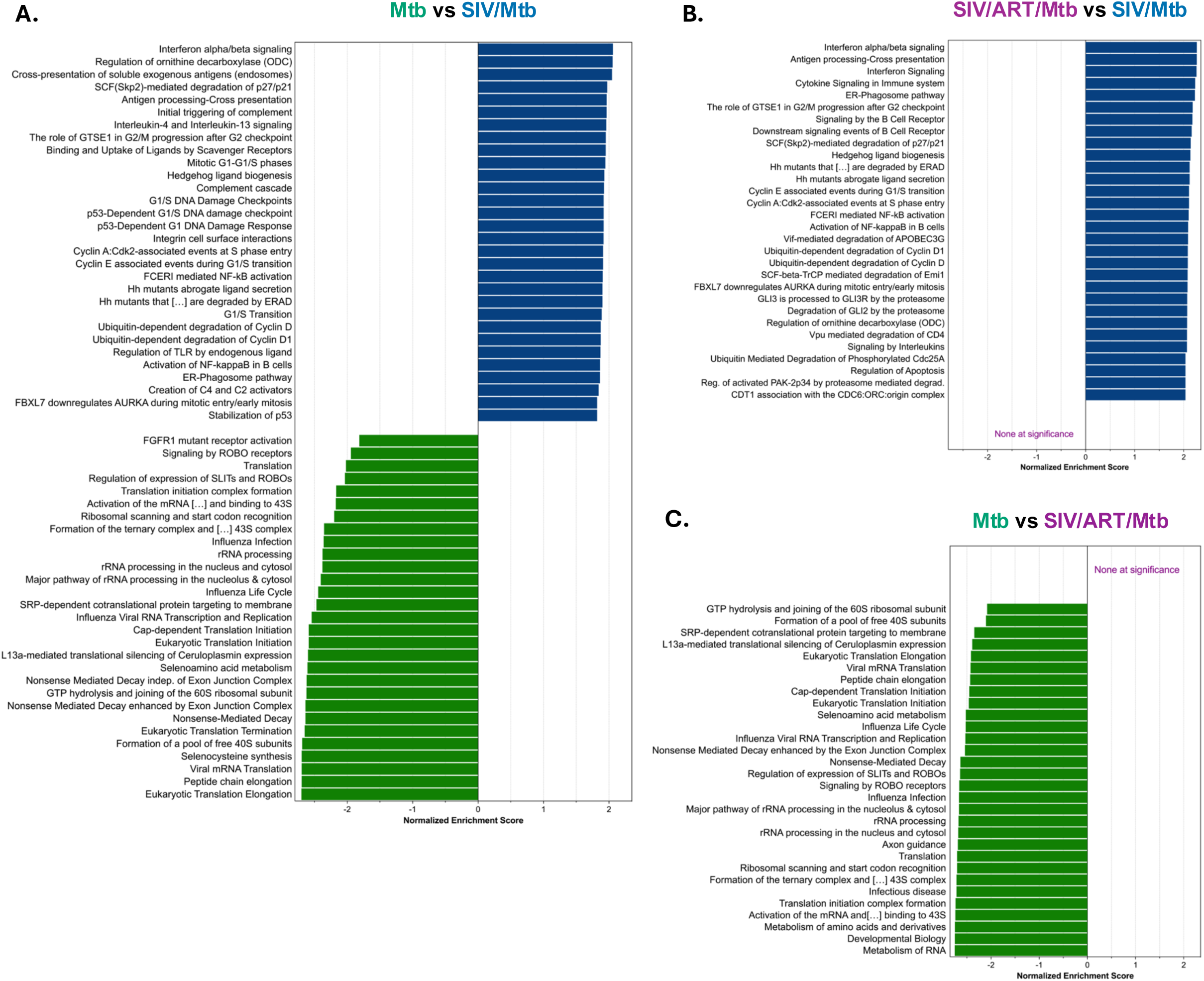
Extended gene set enrichment analysis of myeloid cell function from total granulomas. Full GSEA of all myeloid cells between (A) Mtb vs SIV/Mtb groups (B) SIV/ART/Mtb vs SIV/Mtb groups, and (C) Mtb vs SIV/ART/Mtb groups. Pathways from WebGestalt using Reactome database.

**Supplemental Figure 4.**
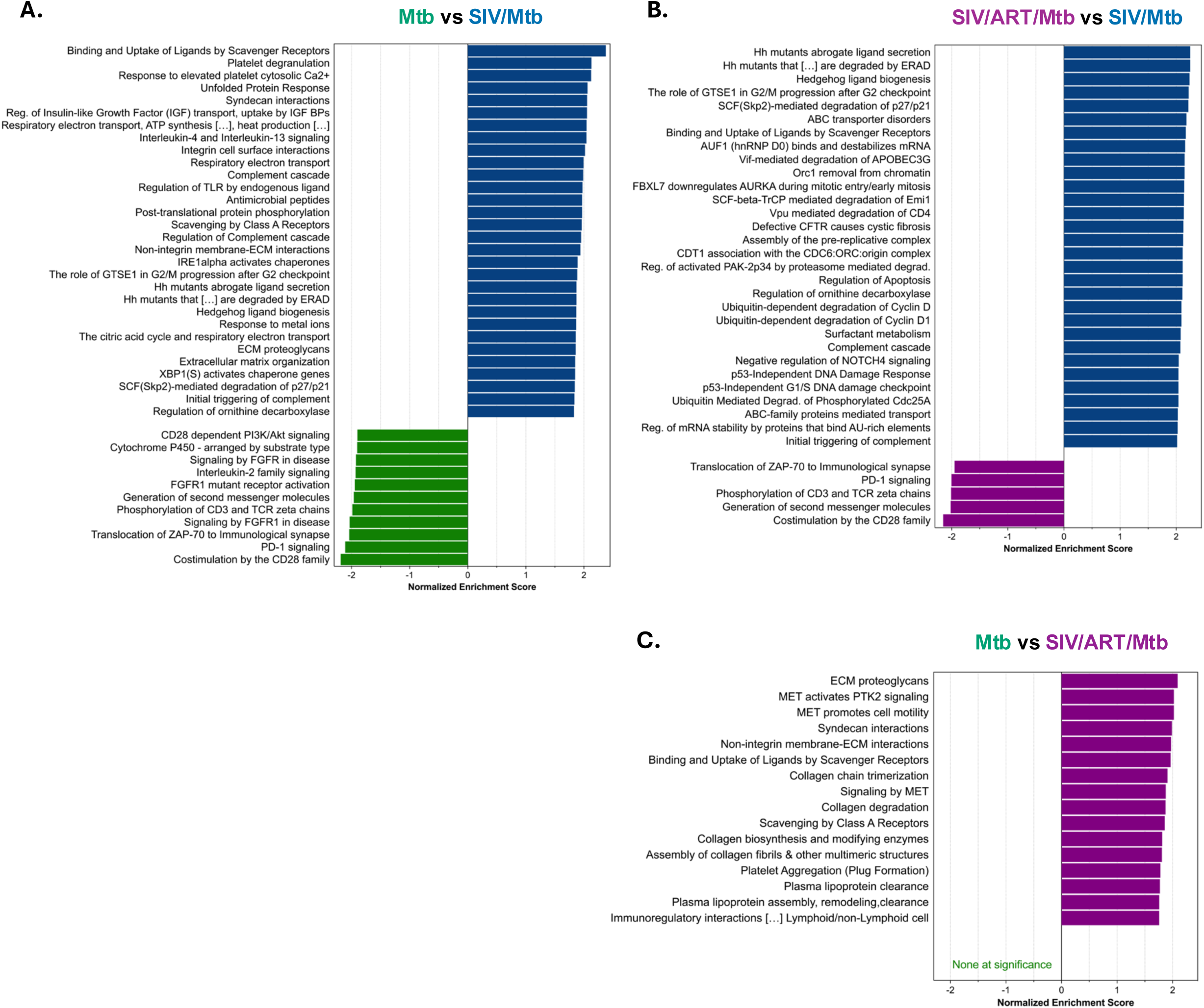
Extended gene set enrichment analysis of T cell function from total granulomas. Full GSEA of all T cells between (A) Mtb vs SIV/Mtb groups (B) SIV/ART/Mtb vs SIV/Mtb groups, and (C) Mtb vs SIV/ART/Mtb groups. Pathways from WebGestalt using Reactome database.

**Supplemental Figure 5.**
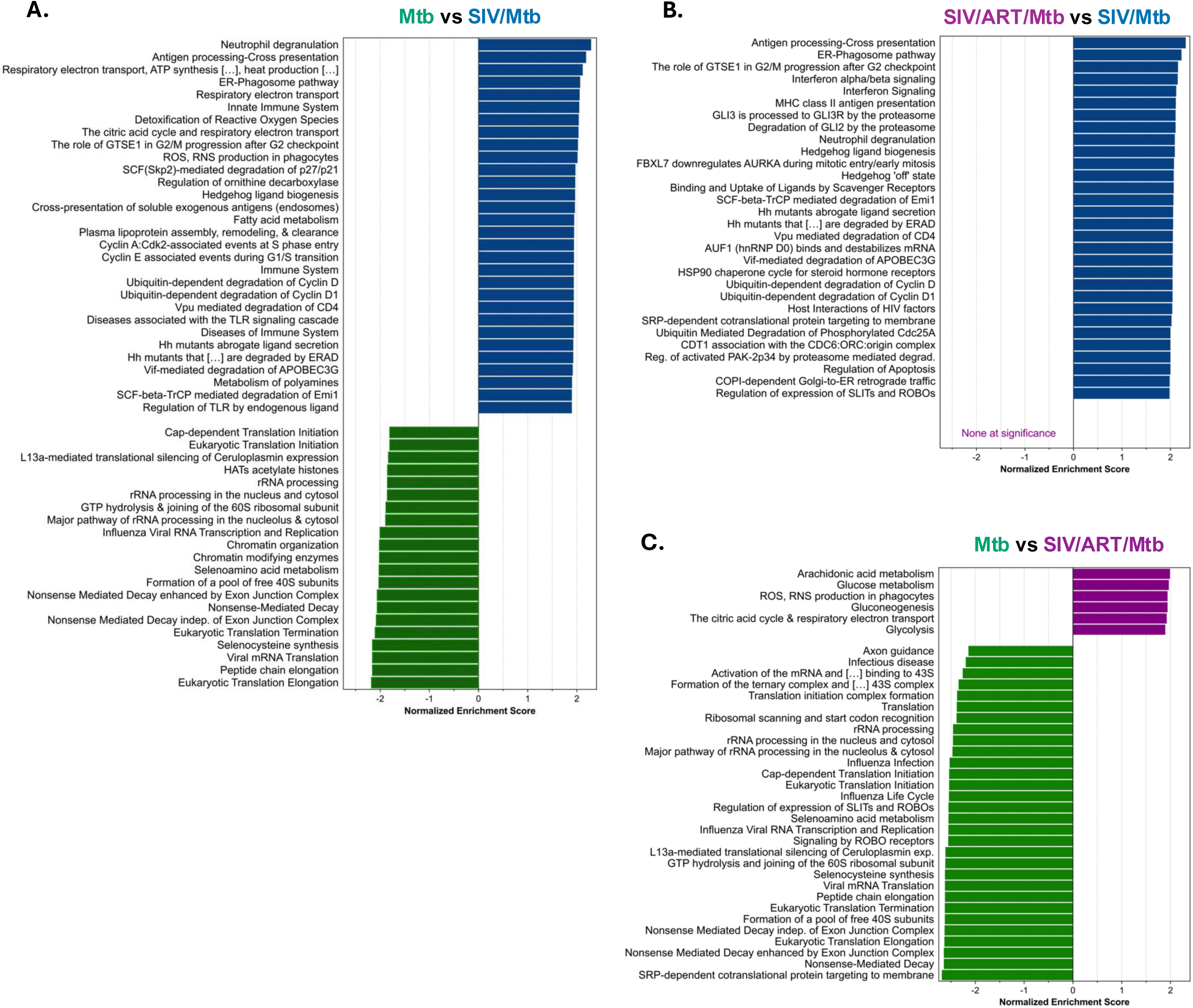
Differential gene expression in myeloid cells in the outer rim of granulomas across groups. Full GSEA of myeloid cells in the outer ring between (A) Mtb vs SIV/Mtb groups (B) SIV/ART/Mtb vs SIV/Mtb groups, and (C) Mtb vs SIV/ART/Mtb groups. Pathways from WebGestalt using Reactome database.

**Supplemental Figure 6.**
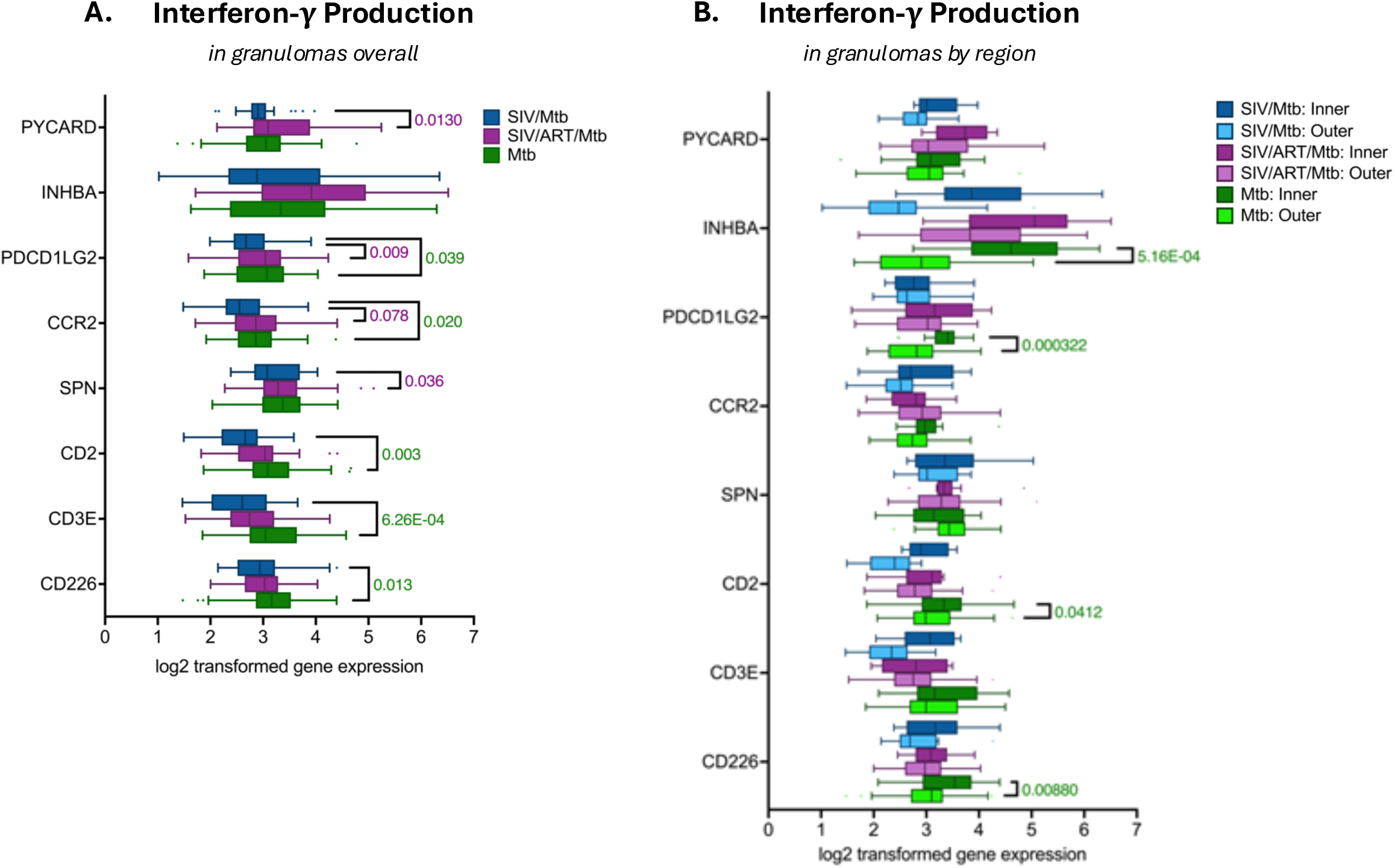
Differential gene expression in T cells in the inner rim of granulomas across groups. Full GSEA of T cells in the inner ring between (A) Mtb vs SIV/Mtb groups (B) SIV/ART/Mtb vs SIV/Mtb groups, and (C) Mtb vs SIV/ART/Mtb groups. Pathways from WebGestalt using Reactome database.

**Supplemental Figure 7.**
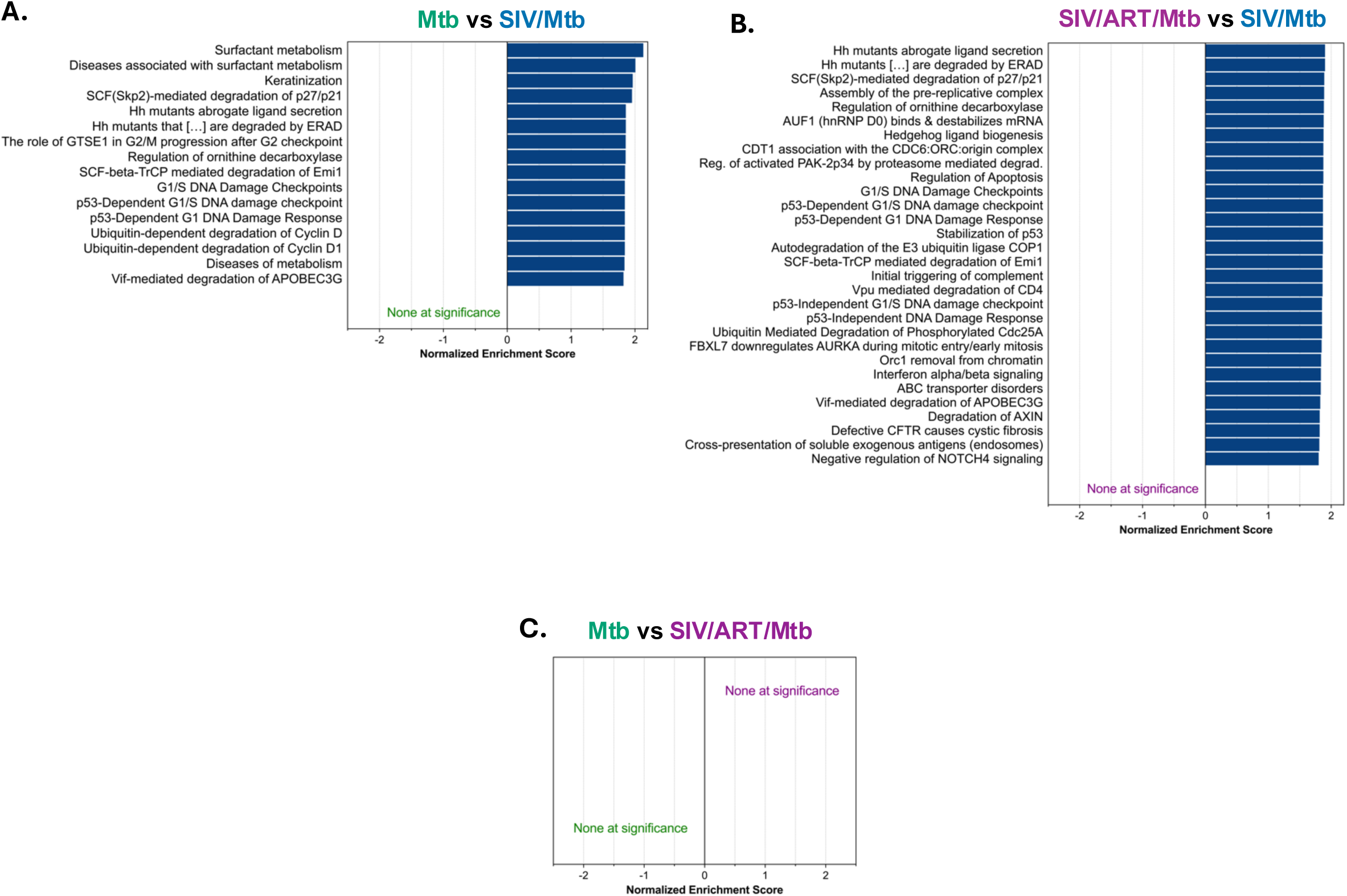
Interferon-γ production in T cells from granulomas. (A) Log2-transformed expression of genes in involved interferon-γ production in all T cells in granulomas. Genes from the GSEA leading edge, only adjusted p-values <0.05 listed. (B) Log2-transformed expression of genes in interferon-γ production in inner versus outer region T cells in granulomas. Genes from the GSEA leading edge, only adjusted p-values <0.05 listed.

**Supplemental figure 8.**
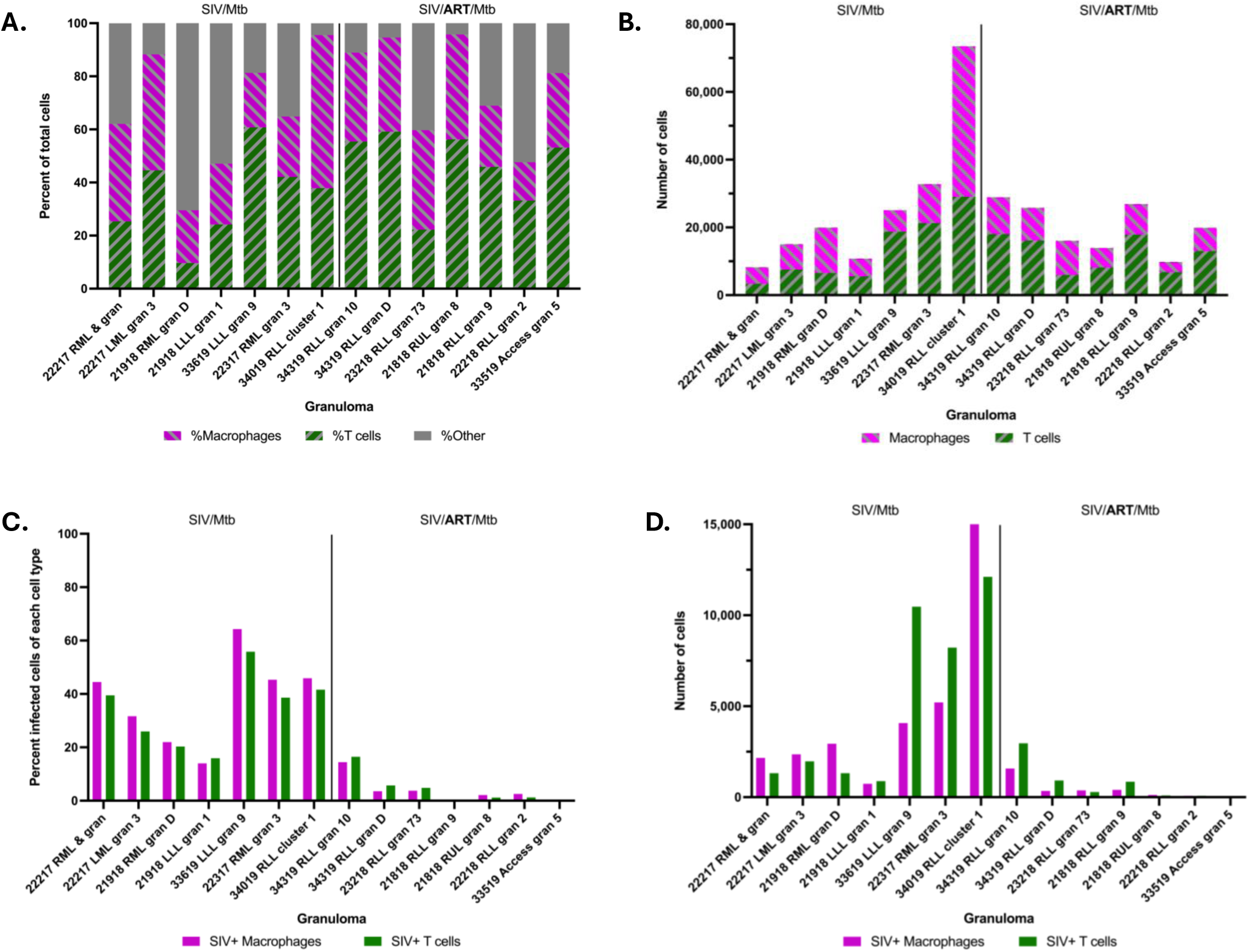
Expanded quantification of SIV-infected cells in granulomas. (A) Percent of CD68+CD11c+ myeloid cells (magenta) and CD3+ T cells (green) of all total DAPI+ cells in granulomas. Grey bars are CD3-CD68-CD11c-DAPI+ cells. (B) Total number of CD68+CD11c+ myeloid cells (magenta) and CD3+T cells (green) in granulomas. (C) Percent of myeloid cells (magenta) and percent of T cells (green) that had detectable SIV RNA in granulomas. (D) Total number of SIV+ myeloid cells (magenta) and SIV+ T cells (green) in granulomas.

**Supplemental Figure 9.**
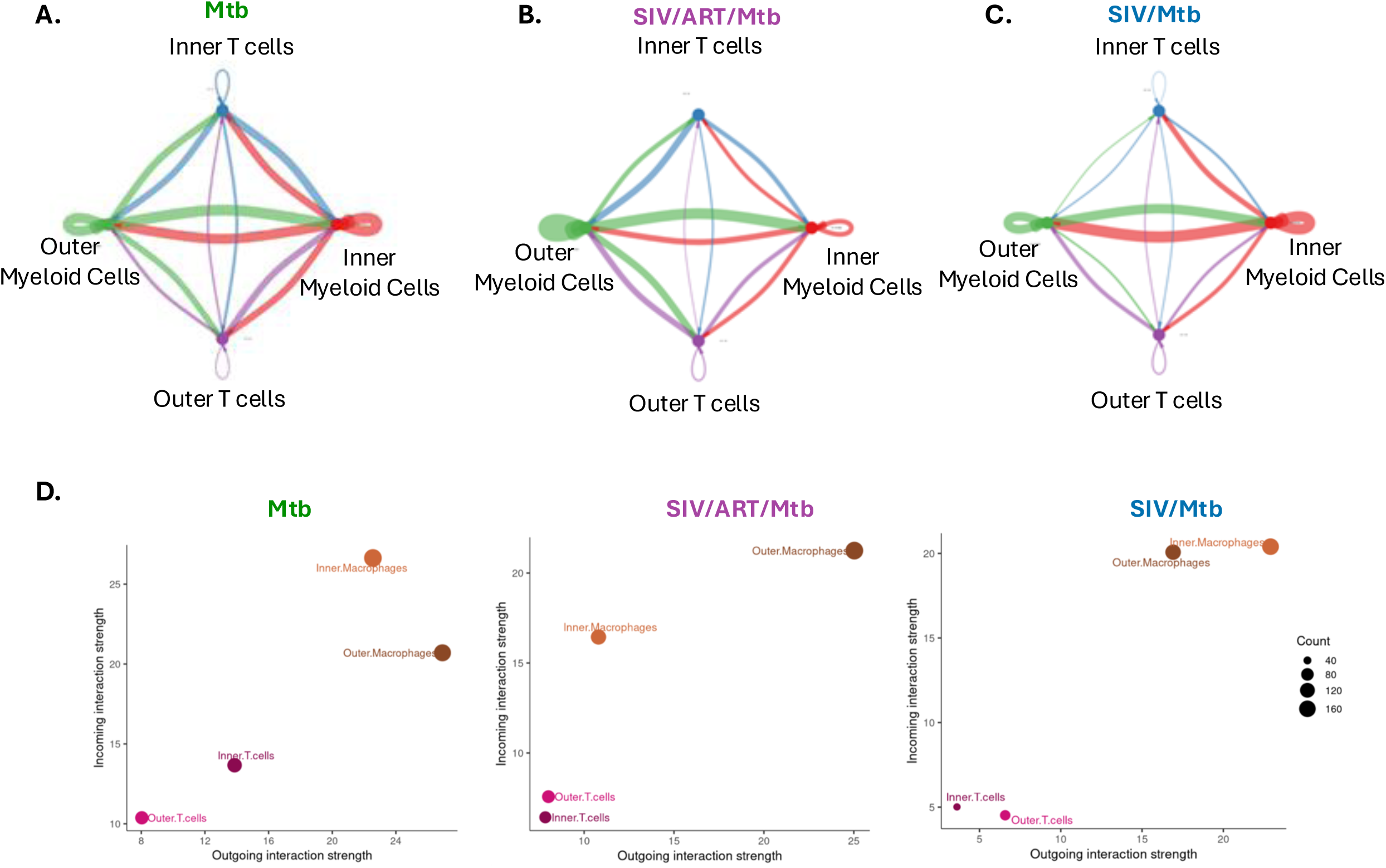
Extended analysis of cell-cell communication in granulomas. (A-C) Aggregated cell-cell communication network between inner and outer myeloid cells and T cells in (A) Mtb granulomas, (B) SIV/ART/Mtb granulomas, and (C) SIV/Mtb groups. Arrows indicate signal senders to signal receivers. Arrow thickness is proportional to the number of ligand-receptor interactions between a given pair. Node size is proportional to the number of ROIs in that cell type. Colors indicate sender group: green is outer macrophages, red is inner macrophages, purple is outer T cells, and blue is inner T cells. (E) Comparison of aggregated incoming and outgoing signal strength in inner and outer myeloid cells and T cells from Mtb (left), SIV/ART/Mtb (middle) and SIV/Mtb (right) groups. Dot size indicates the number of ligand-receptor pairs associated with each group.

